# Containing Cancer with Personalized Minimum Effective Dose

**DOI:** 10.1101/2022.03.28.486150

**Authors:** Masud M A, Jae-Young Kim, Eunjung Kim

## Abstract

Resistance to treatment is a challenge in many cancer therapies. This is partly due to the heterogeneous nature of tumors, where drug-sensitive and drug-resistant cells compete for the same resources. This competition is largely shaped by cancer treatment. The rapid reduction of drug-sensitive cell population during therapy with a maximum-tolerated dose relaxes competitive stress on the drug-resistant cell population, promoting relapse. Therefore, maintaining a high level of drug-sensitive cell population with a treatment break or lower dose can impose effective competitive stress on drug-resistant cell populations. Adaptive therapy (AT) exploits the competition between cancer cells. However, given the heterogeneous treatment response of individual patients, determining a personalized optimal treatment that can fine-tune competitive stress remains challenging. Using a deterministic model of cancer cell population competition, this study defines an effective dose window (EDW) as a range of doses that conserve sufficient sensitive cells, while maintaining the tumor volume below a threshold (e.g., initial tumor volume), to maintain a sustained competition against resistant cells. As a proof of concept, we sought to determine the EDW for a small cohort of patients with melanoma (n=8). We first fitted the model to longitudinal tumor response data from each patient. We performed structural and practical identifiability analyses to confirm the reproducibility and uniqueness of the estimated parameters. Then, we considered a subset of the cohort with uniquely identifiable parameters and estimated patient-specific EDW. We demonstrated that if the dose belongs to the EDW, the tumor volume for each patient could be indefinitely contained either using continuous or AT strategy. Using the optimal control theory, we concluded that the lower bound of the EDW approximates the minimum effective dose (MED) for containing cancer. Taken together, using tumor biomarker data, this study provides a proof of concept that there may exist a patient-specific EDW that keeps the tumor below a threshold (e.g., initial volume) by maintaining sustained competition on resistant cells.

## Introduction

Despite decades of cancer research, drug resistance remains a major hurdle for improving patient outcomes. Genetic, phenotypic, and microenvironmental heterogeneity in most malignant tumors contribute to drug resistance. Currently, a common practice in cancer treatment is to provide a maximum possible dose to kill drug-sensitive cancer cells with tolerable side effects^1^. This maximum tolerated dose (MTD) therapy can rapidly eliminate drug-sensitive cancer cells. However, drug-resistant cells may flourish because of the lack of intra-tumor competition with drug-sensitive cancer cells^2–6^. Several preclinical studies found that the administration of low doses is more effective than MTD in controlling tumor volumes^7, 8^. The successful administration of MTD fractions in early phase trials has shown to improve treatment outcomes^9, 10^. This has inspired the so-called metronomic therapy (MT), which utilizes a one-fixed dosing schedule ranging from one-tenth to one-third of the MTD to all patients^11, 12^. Predicting the treatment dose remains an open problem because of the heterogeneous response between patients. Considering patient-specific tumor evolution, AT strategies have been proposed^13^.

AT is a type of evolutionary therapy that maintains a tolerable level of tumor volume to maintain the competition between drug-sensitive and drug-resistant cells. AT imposes treatment breaks or reduces doses to hamper the growth of resistant cells by leveraging competitive suppression by maintaining sufficient drug-sensitive cells. AT has shown favorable outcomes in both preclinical and clinical settings. AT trial by Zhang et al.^14^ showed a 13.2-24 month delayed tumor progression in prostate cancer. In this study, MTD was applied until the prostate-specific antigen (PSA) level was reduced to 50% of the initial level for each patient and then treatment was held off until the PSA level returned to the initial level. Strobl et al. showed that treatment holidays scheduled after 50% PSA reduction could delay tumor progression for more than 6 months compared to treatment holidays scheduled after reducing PSA to the base level^15^. In an individual base model setting, the group also reported that AT could delay the progression by about 4 months more, when treatment was halted decreasing the PSA level by 30% instead of reducing it by 50 %^16^. Although the above two results are obtained for different model settings and parameter values, both indicate that less aggressive AT may delay progression. Brady-Nicholls et al.^17^ and Kim et al.^18^ also showed that a lower decline from the initial population during the ‘treatment on’ periods could maintain high competitive stress between drug-sensitive and drug-resistant cells, leading to delayed progression. Moreover, Gallaher et al. reported an AT strategy in which the treatment dose was adjusted at four different thresholds with respect to the initial volume based on the tumor response in each patient^2^. To be precise, the authors proposed a four-point dose modulation to maintain the tumor between 110% −50% of the initial volume under the assumption that containing a tumor at a higher volume could delay progression by achieving a more competitive stress on the resistant strain. The first preclinical AT study performed by Gatenby et al. showed that dose modulation was more effective than a treatment holiday strategy in maximizing competitive stress^19^. They showed that an AT involving consecutive “high dose-low dose” windows that contained tumor volume between 80 −120% of the initial volume significantly delayed disease progression in 84% of cases in a breast cancer xenograft model. The benefits of containing tumors at higher volumes have also been theoretically established^20, 21^.

Given that maintaining a sensitive phenotype is required to suppress resistance, one possible way to delay progression is to maintain a tolerable tumor volume rather than trying to eradicate it^22^. How much less dose would be enough to maintain sufficient drug-sensitive cell populations and tumor volume under control below the tolerable level for each patient? To address this question, several studies have employed the optimal control theory. One of the earliest studies addressing the optimal treatment policy subject to drug resistance showed that minimizing the growth rate of resistant cells delayed progression more than minimizing the tumor volume over the treatment period^23^. Recent theoretical studies have emphasized the importance of drug holidays, as is practiced in AT and MT^24, 25^. Cunningham et al. explored the optimal distribution of a constant cumulative dose over a predetermined schedule (to replicate patient clinical visits) to minimize the tumor volume, tumor variance, and total resistant cell density^26^. The virtual patients were divided into three categories according to their response to treatment, as determined by the competition coefficients and initial resistance. Finally, the study recommended delaying treatment as much as possible and administering the smallest possible dose when required, irrespective of the patient group. Moreover, it was shown that if stabilization is possible, an increasing dose titration strategy leads the tumor towards equilibrium^27^. In an in vitro study, a similar dose profile was found to stabilize tumors of two lineages of lung cancer cells^5^. Additionally, Carrere et al.^5^ formulated an optimal control model with singular control to reduce the tumor volume in the same experimental setting. They reported the biologically optimal dose as a periodically increasing dose titration. Recently, a theoretical study by Ledzewicz^28^ and another in vitro study by Bondarenko et al.^29^ also reported a biologically optimal dose^5^ to reduce resistance. However, all existing literature addresses the issue of the optimal dose in theoretical or experimental settings. There are no studies addressing this issue with clinical data. Another issue that draws attention is whether there exists a dose or a range of doses that could effectively contain the tumor. In that case, how is this related to the optimal dose and patient-specific tolerance level?

Further quantitative understanding is urgently required to address these issues. We developed a simple logistic growth model of two different tumor cell populations to explore the dynamics of tumor cell populations. By analyzing the stability of the equilibria, we established the conditions required for containing tumors within a tolerable volume, such as maintaining the initial tumor volume. Our analysis showed that if an equilibrium representative of the tolerable tumor volume exists, administration of a fraction of the MTD belonging to an EDW may redirect the cell population dynamics to the tolerable equilibrium and contain the tumor for a long time. We applied our model and analysis of EDW to a small cohort of melanoma patients whose tumor burden change data was available from a previous study^18^. The fitting of the model to the data generated a set of parameters for each patient. To confirm parameter identifiability, both structural and practical identifiability analyses were performed. We considered a subset of patients whose tumor burden dynamics could be explained using unique parameters. We proposed an optimal control model to characterize the time-dependent MED required to minimize the tumor volume for each patient. Next, we solve the optimal control model with the estimated parameter values which shows that there may exist an optimal dose, continuous administration of a fraction of MTD may direct tumor growth towards drug-sensitive cells only. Further, we simulated different types of AT for each patient by varying the treatment dose and treatment break threshold (i.e., pause level). The time to progression (TTP) of each patient under AT was affected more by the dose level than the pause level. We observed that administration of a dose belonging to EDW resulted in a higher TTP in either the continuous or AT strategy. Most importantly, the optimal dose required to contain cancer at its initial volume is comparable to the lower bound of the EDW, which we defined as the MED.

## Methods

### Mathematical model

We developed an ordinary differential equation model that explains the competition between drug-sensitive and drug-resistant cell populations with logistic growth, as follows:

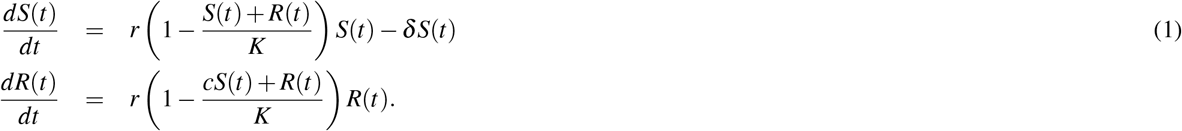

Here, *S*(*t*) and *R*(*t*) denote the populations of sensitive (S-cell) and resistant (R-cell) cells, respectively, at time *t*, and *r* is the intrinsic growth rate of both S and R cells. The term *δ* > 0 is the drug-induced death rate of S cells under treatment. In the absence of therapy, *δ* = 0. The S- and R-cell populations share the same carrying capacity *K*, the maximum size of the tumor owing to resource constraints. Coefficient *c* is a competition coefficient that determines the degree to which the S-cell population inhibits the growth rate of the R-cell population. If *c* < 1, then the S-cell population has a smaller competitive effect on R cells than R cells have on themselves. A coefficient greater than 1 (*c* > 1) implies that S cells have a greater competitive effect on R-cell growth than R cells have on themselves. In this study, we assumed that *c* > 1 based on experimental evidence^2^. The S-nullcline (set of points where 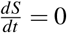) and R-nullcline (set of points where 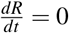) are given by

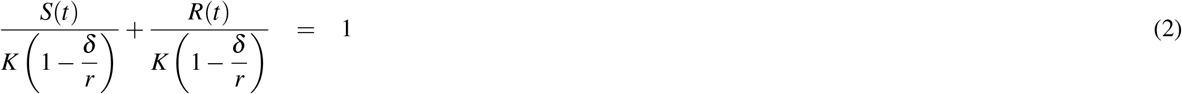

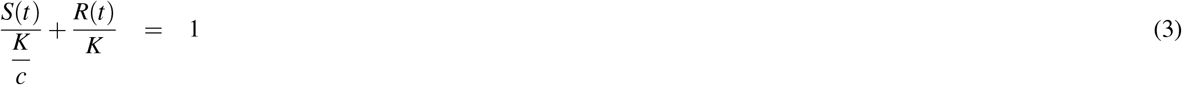

The model exhibits four equilibria, at which both the S-cell and R-cell populations become constant (i.e., 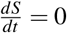 and 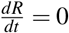). The trivial equilibrium is (*S, R*) = (0, 0) and the S-only equilibrium is 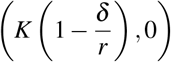. R-only equilibrium (0, *K*). The coexistence equilibrium is 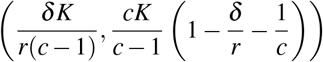,

### Parameter identifiability and model parameterization

Parameter identifiability analysis assesses how well the parameters of a model can be estimated by experimental or clinical data, assuming that a mathematical model fits the data well with a small error. The goodness of fit does not guarantee the reliability of the estimated parameters. For instance, low-quality data with high noise or a small number of data points may result in various parameters that can fit the data almost equally well. Identifiability analysis has become more important, particularly in modeling biological systems with often partially observed noisy data^30–32^. First, we performed a structural identifiability analysis to assess the inherent properties of the model. Next, we fit the model to the data and assessed the practical identifiability of the estimated model parameters.

### Structural identifiability

Structural identifiability is an inherent property of a model that addresses the existence of a unique set of parameter values given noise-free observations at all-time points^33^. To formally analyze the identifiability, we rewrote the system (1) as

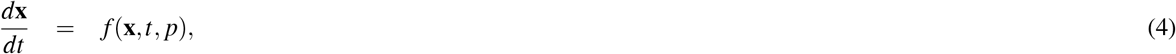

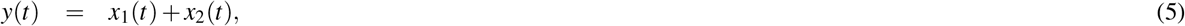

where the output *y*(*t*) denotes the data (i.e., the tumor burden at time *t*), **x** is the state vector of *S* and *R* (**x** = [*S, R*]), and **p** indicates a vector of unknown parameters (**p** = [*r, K, δ, c*]). For models (4) and (5), the parameter *p*_*i*_ in **p** is uniquely or globally structurally identifiable if for almost all initial conditions and almost every **p**\*,

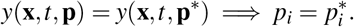

The above equation resembles a one-to-one relation between the output and the parameters. This can be rephrased as the injectivity of the map *ϕ* : **p** → *y*, defining the model output as a function of the parameters **p**^34^. We adopted a differential algebra approach^34–36^ to address the structural identifiability of **p**, which is summarized in the following steps.

Step 1 Rewrite the model in terms of the output *y* and parameters **p** to express the dependency of the observable on the parameters. This is known as the input-output equation^37^.
Step 2 Normalize the input-output equation by the coefficient of the highest ranking monomial of *y* to deduce the monic polynomial^38^.
Step 3 Examine the injectivity of the coefficients of the monic polynomial with respect to the parameters, which confirms the structural identifiability of the model^36^.

This approach can be implemented using the free web-application COMBOS^33^, which has been used in several previous studies to assess structural identifiability^39–41^. Therefore, we chose to use the COMBOS^33^ web application to verify the structural identifiability.

### Parameter estimation

We employed the maximum-likelihood approach to estimate the parameters. We assumed that the tumor burden *V* (*t*_*n*_) at time *t*_*n*_ is a sample from the Poisson distribution with mean *y*(*t*_*n*_; **p**). Using the probability mass function of the Poisson distribution, we derived the likelihood of observing the longitudinal tumor burdens *V*(*t*_1_),*V*(*t*_2_), …, *V*(*t*_*N*_) at times *t*_1_, *t*_2_, …*t*_*N*_, as follows:

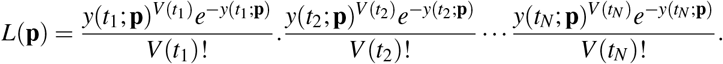

Next, we formulated negative log likelihood (*NLL*) as follows

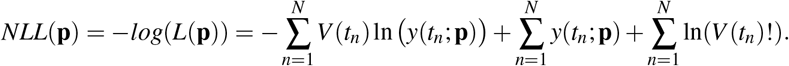

An optimization algorithm was employed to identify the parameters that minimized the above equation (minimizing the likelihood). In this study, we employed the Nelder-Mead Simplex method built into the MATLAB function *fminsearch*. It is worth mentioning that, by minimizing *NLL*, we maximized the probability of realizing the observed data.

### Practical identifiability

Practical identifiability concerns the quantity of data required to determine parameters, whether given the amount of data, one can uniquely infer the parameter values. The analysis was performed locally by perturbing the estimated parameters to fit the data. Specifically, we utilized the Fisher information matrix (FIM) and profile likelihood (PL) approaches.

To derive an FIM, we first calculated a sensitivity matrix **M**,

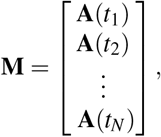

where **A**(*t*_*n*_) is an *n*_*x*_ × *n*_*p*_ matrix (*n*_*x*_ is the number of state variables and *n*_*p*_ is the number of parameters). An element of **A**(*t*_*n*_) is defined by 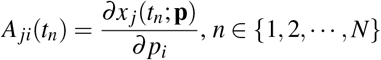.

FIM is defined as **F** = **M**^*T*^ **M**, and its rank indicates the number of identifiable parameters. A rank of *n*_*p*_ indicates that the number of parameters *n*_*p*_ is practically identifiable^42, 43^. A finite-difference method was applied to approximate **F**. We perturbed each 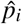 to 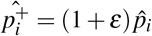 and 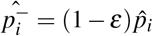, where *ε* = 0.001. For this, we simulated the model and numerically approximated the derivatives 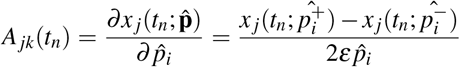. Note that 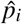 resembles the estimated value of *p*_*i*_. Moreover, we computed the profile likelihood for a parameter *p*_*i*_ by varying the parameter over an interval containing 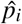 and fitting the remaining parameters^44^. The resulting likelihood for each *p*_*i*_ constitutes the profile-likelihood function for *p*_*i*_. Mathematically, it can be written as

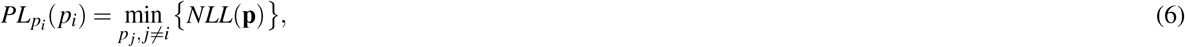

Where 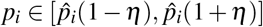, and *η* = 0.2. If all profile likelihoods show a global minimum at the estimated value of the parameters, then the parameters are practically identifiable.

### Optimal control

To optimize the drug dose, the following optimal control process was employed: We multiplied the time-dependent dose modulation parameter *u*(*t*) ∈ [0, 1] by *δ* in model (1) and obtain the resulting model (7).

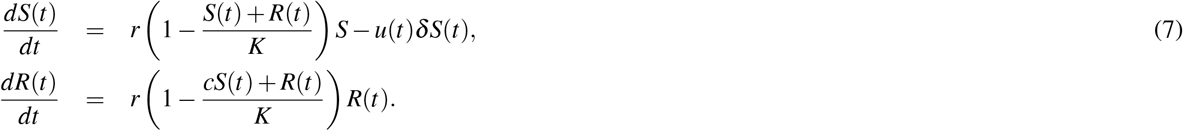

Here, control parameter *u*(*t*) denotes the required optimal dose as a fraction of the MTD. In the optimal control problem, we aimed to minimize the tumor volume using the possible minimum fraction of MTD. Following this aim, we model the cost functional (8)

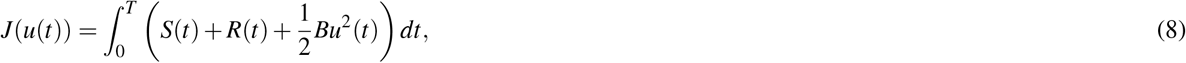

where *B* is the cost or weight of the dose. Our objective is to find *u*\*(*t*) such that

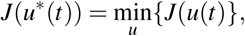

where *u*(*t*) is a measurable function, and 0 ≤ *u*(*t*) ≤1. We considered the quadratic form of the control term to deduce the time-dependent continuous dose. The convexity of the integrand of the cost functional (8) implies the existence of an optimal dose *u*\*(*t*)^45^. We used Pontryagin’s maximum principle^46^ to derive the necessary conditions for the optimal solution, defined the Hamiltonian (ℋ) in equation (9) from which we derived the adjoint system (10), and characterized the time-dependent optimal control in equation (11) (see Theorem C.1 for details).

Hamiltonian,

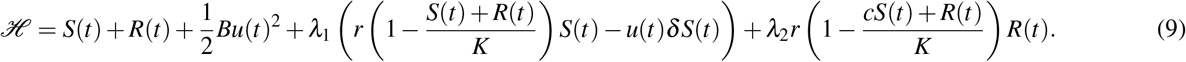

Adjoint system,

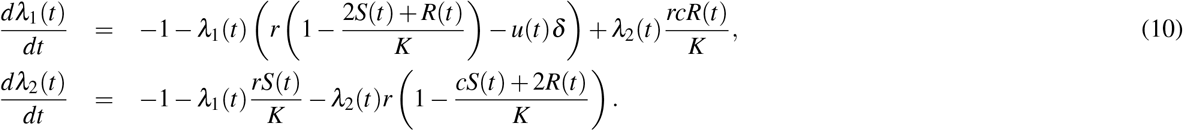

Time dependent Control,

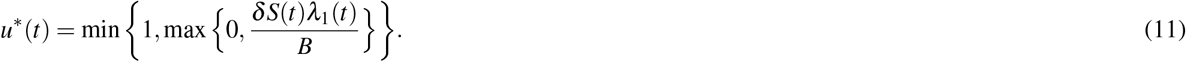

The forward–backward sweep method^47^ is used to numerically solve the optimality system consisting of equations (7), (10), and (11).

### Adaptive therapy

We considered various doses and pausing levels of adaptive therapy, defined as follows.

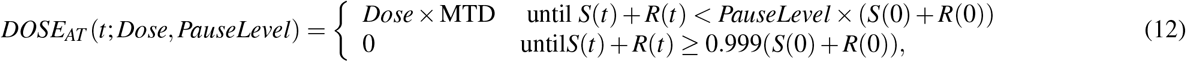

Where *Dose* and *PauseLevel* ∈[0, 1]. We simulated the treatment until the tumor volume reached the pause level relative to the initial volume, and held the treatment off until the tumor burden increased up to 99.9% of the initial burden.

## Results

Following the mathematical model analysis described in the Methods section, we first defined the EDW. We then applied the model to eight melanoma patients to derive patient-specific EDW and compared treatment outcomes under both continuous and AT with different doses and pause levels. Finally, by employing the optimal control theory, we derived a patient-specific MED that can indefinitely control tumor volume.

### EDW

Mathematical analysis of the model (1) shows that the dynamics precisely depend on three model parameters:intrinsic growth rate of S cells (*r*), drug-induced death rate (*δ*), and competition coefficient (*c*) of S cells over R cells (please refer to Appendix A). The dynamics can be classified into the following three categories and are graphically represented in Fig 1.

**Figure 1.**
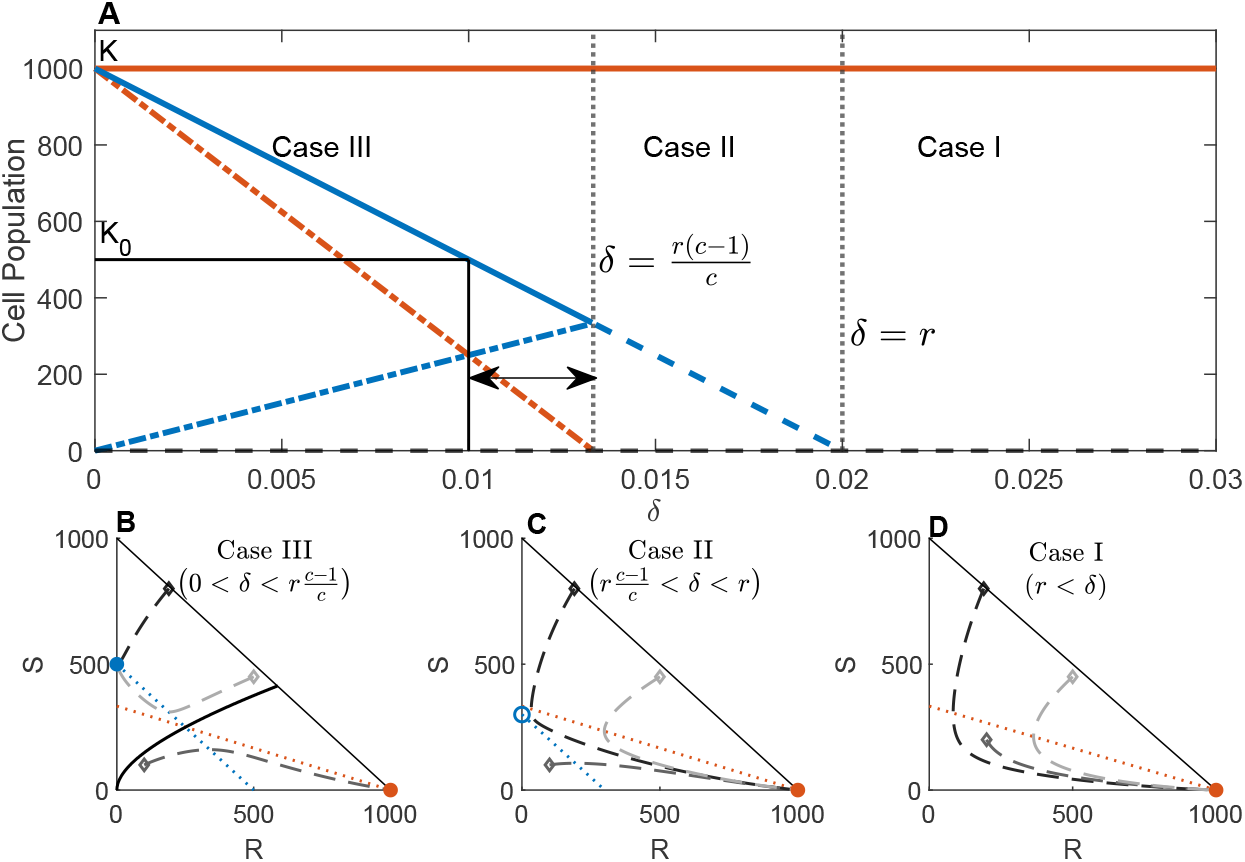
The upper panel (A) shows the bifurcation diagram. *K*_0_ is the initial tumor volume, which is assumed to be the threshold tumor burden that determines tumor progression. The vertical grey dotted lines divide the domain into three parts, showing the equilibria for Cases I, II, and III. The solid orange line shows the stable R-only equilibrium and the solid blue line shows the stable S-only equilibrium. The dashed blue line indicates an unstable S-only equilibrium. The dash-dot blue and orange lines indicate the S-cell and R-cell populations, respectively, in the unstable coexistence equilibrium. The solid black line represents the initial cell population and corresponding drug-induced death rate. The horizontal double arrow indicates the EDW. The lower panel shows the phase diagrams for cases III (B), II (C), and I (D) (from left to right). Triangular regions indicate the phase space. The dotted blue and orange lines represent the S- and R-nullclines, respectively. The dashed lines with different shades of gray are the trajectories starting from different points in the phase space. The solid black curve (B) shows the separatrix in case III. The orange filled dots show the stable R-only equilibrium in all the cases. The blue empty dot shows the unstable S-only equilibrium in Case II, and the blue filled dot in Case III shows a stable S-only equilibrium. It is observed that the trajectories starting from the same three points evolve in a different manner as *δ* changes. The assumed parameter values for the above diagram are *r* = 0.02, *c* = 3, *K* = 1000, and *K*_0_ = 500, and the initial conditions for the phase portraits are (*S*(0), *R*(0)) = (800, 190), (200, 200), and (450, 500).

- Case I (*δ > r*): The system exhibits monostable dynamics with an unstable trivial equilibrium (*S, R*) = (0, 0) shown by the dashed black line and a stable R-only equilibrium (*S, R*) = (0, *K*) shown by the solid orange line in Fig 1A and orange filled dot in Fig 1D (inferred from Theorem A.1 and A.3). An S-nullcline did not exist in this case. All example phase portraits converge to the R-only equilibrium.
- Case II 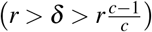: The system exhibits monostable dynamics with an S-only unstable equilibrium 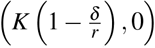 (shown by dashed blue line in Fig 1A and blue empty dot in Fig 1C) in addition to the above two equilibria (inferred from Theorem A.1,A.2 and A.3). Although both nullclines exist, they do not intersect, and hence, no coexistence equilibrium exists. In this case, all phase portraits converge to the R-only equilibrium.
- Case III 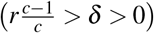: In this case, as *δ* goes below the threshold 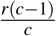 the null clines intersect at the coexistence equilibrium 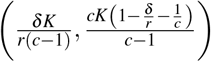 shown by the dotted blue (S cells) and orange (R cells) lines in Fig 1A, which is unstable (Theorem A.4). Concurrently, the S-only equilibrium becomes locally asymptotically stable (shown by the solid blue line in Fig 1A and the blue-filled dot in Fig 1 B). As a result, the system exhibits bistable dynamics with locally asymptotically stable S-only and R-only equilibria and unstable coexistent equilibrium (inferred from Theorems A.1, A.2, A.3, and A.4). The coexistence equilibrium and trivial equilibrium lie on the separatrix of the basin of attraction of the two locally asymptotically stable equilibria. Recall that the set of points (i.e., initial condition) starting from which the trajectories converge to an equilibrium is the basin of attraction of the equilibrium. The separating boundary between the basins of attraction of the two equilibria is the separatrix. In Fig 1B, the solid black curve is the separatrix that partitions the interior of the phase space into the basin of attractions of the two stable equilibria (S-only equilibrium: 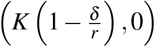, and R-only equilibrium: (0, *K*)). We observed that the trajectories starting above and below the solid black line (separatrix) converge to the S-only 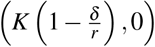 and R-only ((0, *K*)) equilibria, respectively.

In cases I and II, the drug-induced death rate was too high for all S-cells to compete against R-cells. Consequently, S-cells die out and the total cell population approaches the R-only equilibrium. In these cases, the cancer cells grow to their carrying capacity. In case III, the drug dose making 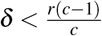 can maintain a sufficient number of S cells to win the competition against R cells and suppress their growth. As a result, R-cells die out and the total cell population approaches an S-only equilibrium, provided that the initial cell combination belongs to the basin of attraction of S-only equilibrium. Again, the coexistence equilibrium lies on the separatrix, which depends on the drug-induced death rate *δ*. Thus, by modulating the drug dose, the basin of attraction of the S-only equilibrium can be expanded, and the dynamics can be directed towards the S-only equilibrium (see an example case in Section **Effect of dose modulation on tumor cell population dynamics**). If the trajectory approaches the S-only equilibrium 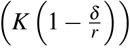, the cancer cells only grow to a level below the carrying capacity under therapy. In this study, we named this level as tolerable tumor volume. In the clinic, Response Evaluation Criteria in Solid Tumor (RECIST), version 1.1 are used to evaluate patients’ response to cancer therapy^48^. In RECIST, if the sum of the diameters of target clinical lesions increases by at least 20% from the initial sum before therapy, the disease is called a progressive disease. If the sum increases by less than 20% and decreases by less than 30% (70% < tumor volume < 120% from the initial), it is called a stable disease. One can consider the tumor volume in stable disease as a tolerable volume. In this study, we assumed that the initial tumor burden (*K*_0_) was the tolerable tumor volume. Thus, if the S-only equilibrium is smaller than the initial tumor burden 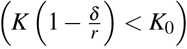 in Case III 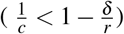, it can be claimed that the cancer can be contained at a tolerable level. Combining these results, we found that successful containment requires a dose that satisfies equation (13).

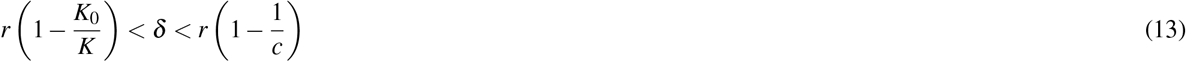

We defined equation (13) as the EDW, which is indicated by the horizontal double arrow in Fig 1A. Our analysis shows a suitable fraction of MTD belonging to EDW (rather than MTD) could be more effective in containing cancer cell population growth. The upper bound of EDW depends on the growth rate *r* and the competition coefficient *c*. As *c* increases, the upper bound approaches *r* (as 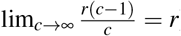); however, the sensitivity of the upper bound to *c* is very high when *c* is slightly above 1 (as 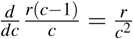 is a decreasing function for *c* ≥ 1). Therefore, slight competitive stress of the S cell over the R cell may offer suitable EDW. Additionally, the lower bound of EDW depends on the tolerance level of the patient. Increased levels of tolerable tumor volume decrease the lower bound of the EDW. To relate these findings to real-life scenarios, we fitted the model (1) with the biomarker level data of melanoma patients treated with MTD therapy.

### Patient specific EDW

As treatment response dynamics vary among patients, we expected the EDW to be patient-specific. To demonstrate how one can estimate patient-specific EDW, we applied our model to eight patients with advanced metastatic melanoma. All patients were treated with continuous MTD but showed disease progression within six months of treatment. Melanoma does not have an ideal biomarker for assessing tumor burden. LDH (Lactate dehydrogenase) is clinically used in melanoma to monitor the tumor burden and make treatment decisions^49, 50^. Previously reported temporal dynamics of patient LDH data were employed in this study^18^. For simplicity, we assumed that the LDH level is equivalent to the total number of cancer cells (*S*(*t*) + *R*(*t*)), and during the initiation of treatment, 1% of cells are resistant to all eight patients^51, 52^.

Following the structural identifiability analysis, we determined that a unique set of parameter values, that fit the model to the data, exists. Then, we estimated the model parameters that fit the patient data (Fig 2) by employing the maximum likelihood method described in the section **Parameter estimation**. To assess the practical identifiability, we calculated the rank of the corresponding FIM. It is noteworthy that if the parameters are practically identifiable, the rank of the corresponding FIM is full (i.e., the same as the number of parameters). Because the model has four parameters, practical identifiability requires the rank of the FIM to be four. The analysis revealed that although the data fitting appears reasonable for all eight patient cases, the rank of FIM is the same as the number of parameters for only five patients (Patients 2, 4, 6, 7, and 8). Moreover, we checked whether the profile likelihood 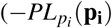 defined in equation (6)) of the patients has a global minimum. The profile likelihoods of the parameters for patients 2, 6, 7, and 8 (Fig. 3) revealed the existence of a unique minimum. For the parameter estimates for patient 4, although FIM has a full rank of 4, practical identifiability was not proved through the profile likelihood (row 3 of Fig 11). Thus, we conclude that the parameters of patients 2, 6, 7, and 8 were identifiable. Table 1 lists the estimated patient-specific parameters. The parameter values vary significantly from patient to patient, which emphasizes the requirement for a patient-specific treatment design. Using the estimated parameters, we predicted the lower and upper bounds of the EDW (columns 6 and 7 in Table 1) following the method described above.

**Table 1.** Estimated values of the identifiable parameters (from the second column to fifth column). The last two columns show the lower and upper bounds of the EDW.

| | $r$ | $K$ | $\delta$ | $c$ | $r\left(1 - \frac{K_0}{K}\right)$ | $r\left(1 - \frac{1}{c}\right)$, |
| --- | --- | --- | --- | --- | --- | --- |
| Patient 2 | 0.1318 | 580.8725 | 0.0890 | 2.2920 | 0.0503 | 0.0743 |
| Patient 6 | 0.1815 | 798.6885 | 0.1376 | 3.1830 | 0.0276 | 0.1245 |
| Patient 7 | 0.1829 | 663.0487 | 0.1283 | 2.3628 | 0.1040 | 0.1055 |
| Patient 8 | 0.1655 | 544.2614 | 0.0968 | 1.7348 | 0.001 | 0.0701 |

**Figure 2.**
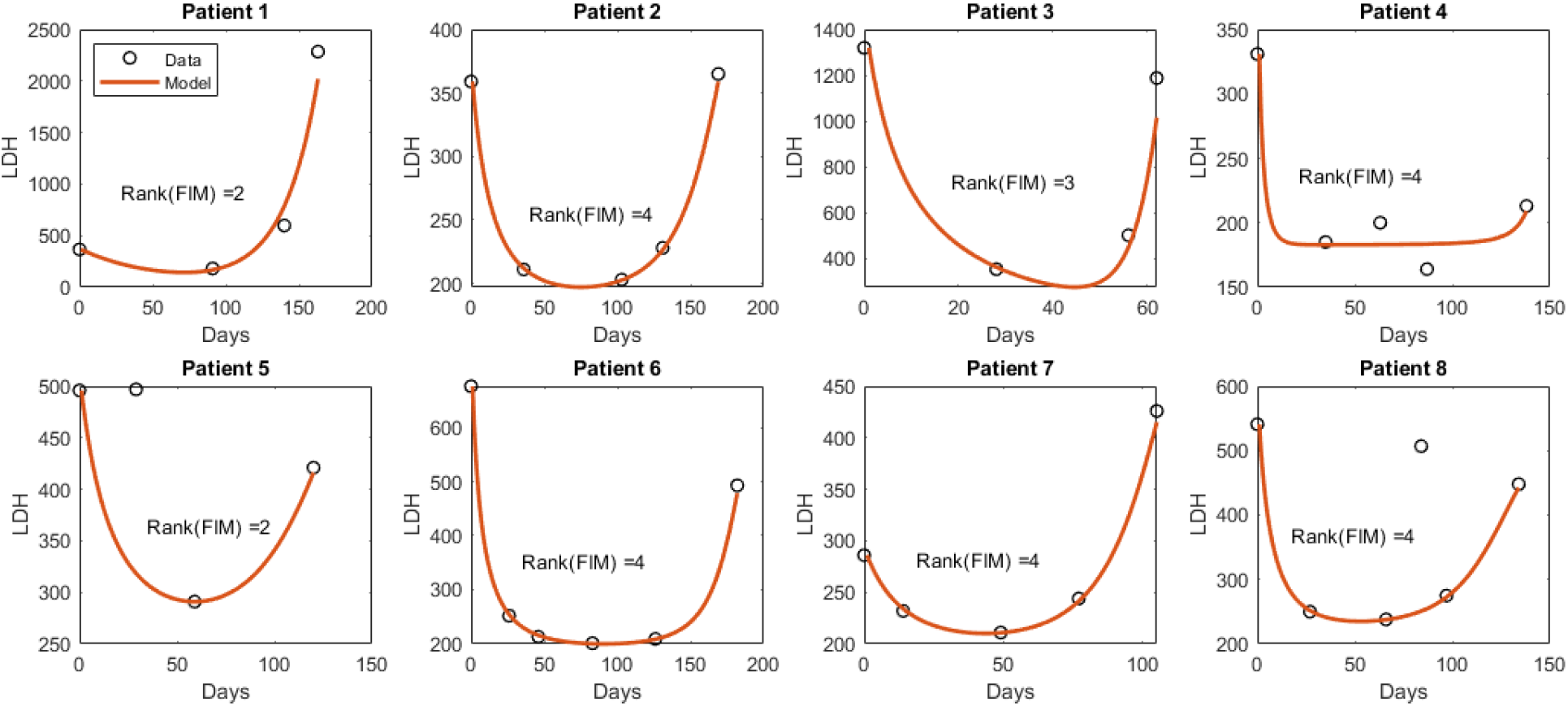
The circles indicate the data points for each patient and the solid line shows the model predicted dynamics of the LDH level. FIM: Fisher information matrix. It should be noted that the second and fourth data points in the case of Patients five and eight, respectively, were excluded while fitting, as these two instances resemble deviations from the regular trend observed through the other data points, which could be a consequence of other physical problems. Owing to the lack of detail in the patient’s history, we proceed with this assumption.

**Figure 3.**
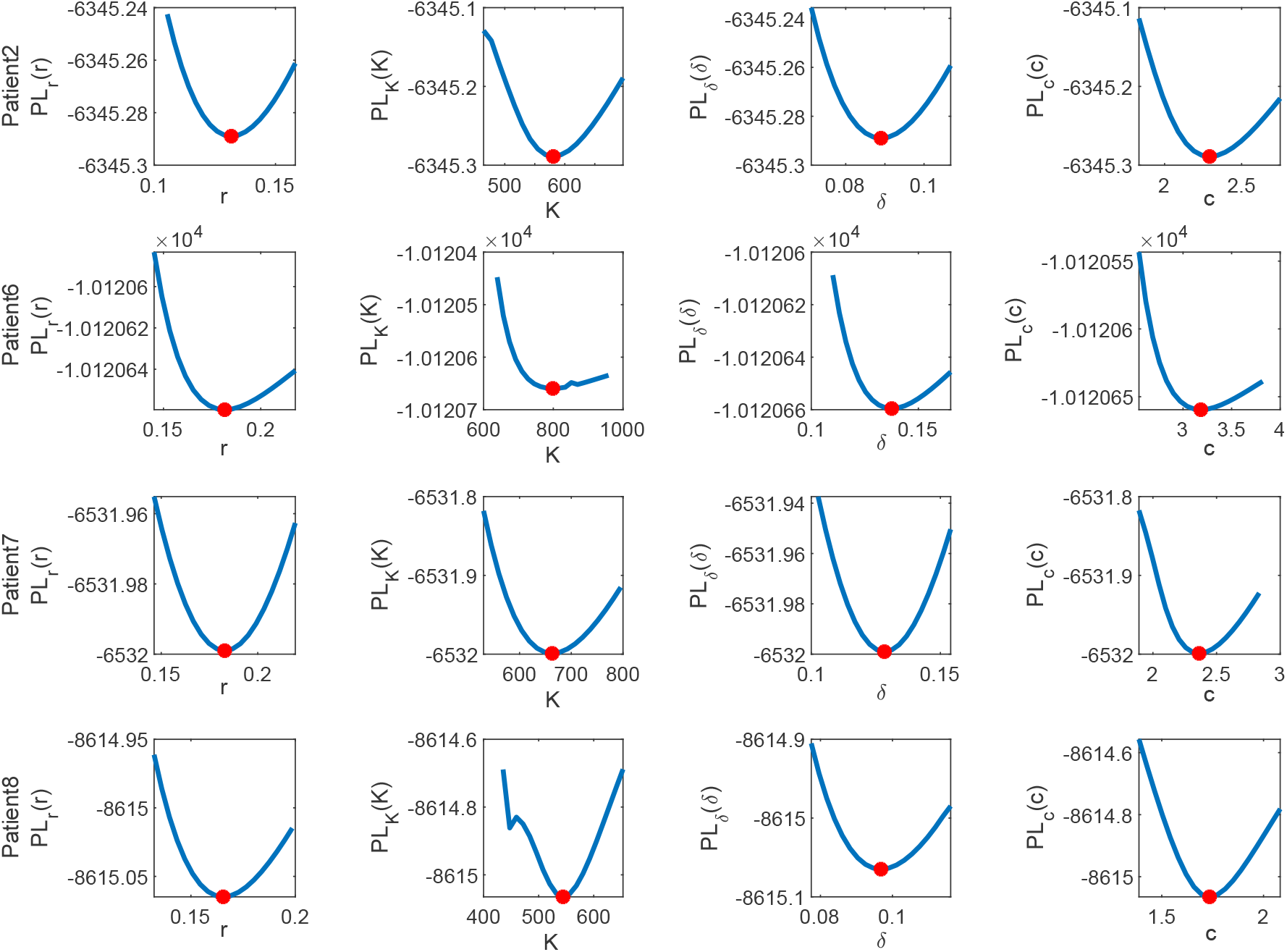
Each row shows the profile likelihood for each of the parameters for Patient 2, 6, 7 and 8 respectively, where y axis is 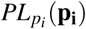 defined in equation (6) and x-axis is the perturbed parameter *p*_*i*_ ∈ {*r, K, δ, c* }. All the curves have minima at the estimated values shown by the red dot, which confirms practical identifiability.

### Effect of dose modulation on tumor cell population dynamics

As discussed in the derivation of the EDW in the Subsection **EDW**, the parameter *δ* plays an important role in determining the intra-tumor composition. For all four patients, the estimated parameter *δ* is greater than the upper bound of the EDW (Table 1, 4th column vs. last column), but less than the growth rate (Table 1, 4th column vs. 2nd column). Therefore, the cell population dynamics belong to Case II for all patients (Fig. 1C), and the R-only equilibrium is the only stable equilibrium that exists under MTD. For all four patients, the cell population dynamics approached the R-only equilibrium and the tumor eventually relapsed despite a significant initial reduction in tumor volume following treatment initiation. Reducing *δ* using a fraction of MTD (smaller *δ* within EDW) could steer tumor dynamics to a more favorable outcome, sensitive cell-only equilibrium, given that the initial composition ((*S*(0), *R*(0))) belongs to the respective basin of attraction. Because we do not have an explicit characterization of the separatrix, we cannot yet determine if the initial condition of S and R cells for each patient belongs to the basin of attraction of S-cell-only equilibrium. However, the coexistence equilibrium, which depends on the drug dose (*δ*), depends on separation. Moreover, the S-nullcline and, hence, the S-only equilibrium, depends on the drug dose (equation (2)). Therefore, the drug dose can be modulated to expand the basin of attraction of the S-only equilibrium to contain the initial point and reach a suitable S-only equilibrium (tolerable tumor burden, e.g., initial tumor volume).

To further illustrate this, we presented a scenario for Patient 2 (Fig 4). The R-nullcline (3), shown by the orange solid line, is invariant to *δ*. The S-nullcline is indicated by a dotted blue line for the MTD. The nullclines do not intersect, and the only stable equilibrium is the R-only equilibrium (filled orange circles). Therefore, the trajectory starting from the initial point (gray diamond (◊)) approaches the R-only equilibrium (filled orange circle), as shown in Fig 1C (Case II). For a dose in the EDW (13) (for instance, 70% of MTD), the S-nullcline (purple dotted line) and R-nullcline intersect at the unstable coexistence equilibrium (the green asterisk (*)), which lies on the separatrix. Trajectories starting above and below the separatrix approach the S-only (purple circle) and R-only equilibrium (orange circle), respectively. Because the initial S-R cell combination (◊) belongs to the basin of attraction of the S-only equilibrium, the cell population approaches the S-only equilibrium (purple circle) (as we observed in Fig 1B (Case III)). When the population reaches S-only equilibrium, it remains at a constant level. Overall, if the dose is modulated so that the drug-induced death rate belongs to EDW, continuous therapy with the modulated dose (fraction of MTD) can contain the tumor at an S-only equilibrium indefinitely.

**Figure 4.**
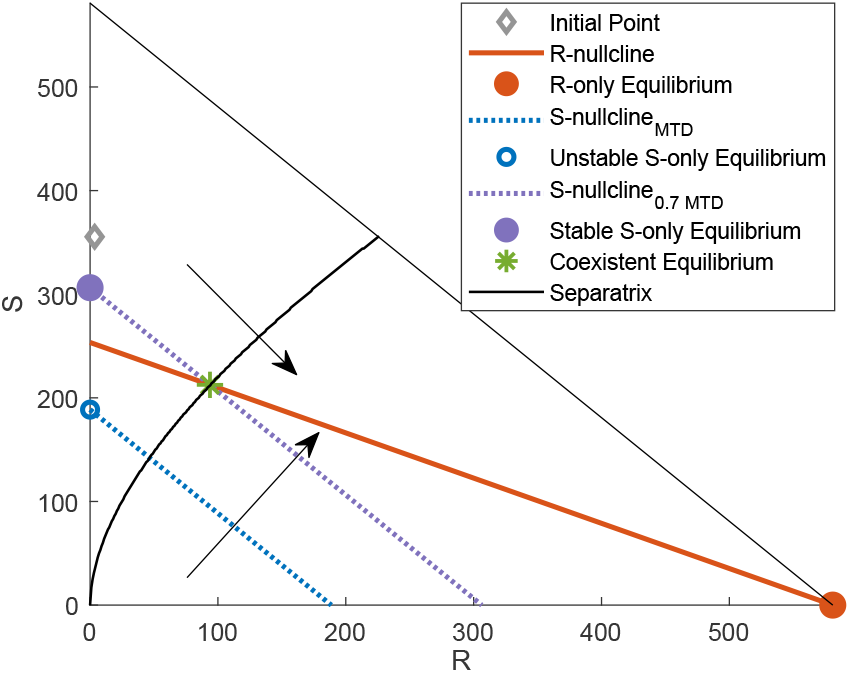
The triangular region shows the phase space for patient 2. The initial cell composition is shown by the grey diamond (◊). The solid orange line shows the R-nullcline (equation (3)), which is invariant to *δ*, and the solid orange circle represents the R-only equilibrium. The dotted blue line shows the S-nullcline (equation (2)) with the MTD, and the empty blue circle is the unstable S-only equilibrium. The dotted purple line shows the S-nullcline (2) with 70% of the MTD (belonging to the EDW ((13))), and the filled purple circle is the stable S-only equilibrium. The solid black line indicates the separatrix. The two arrows indicate the direction in which the separatrix and S-nullcline are shifted when *δ* decreases.

### Dose optimization

We have demonstrated that there exists a patient-specific EDW that can contain a tumor below a threshold. Next, we investigated which dose in the EDW was optimal. To this end, we applied the optimal control theory to derive the optimal dose that contains the cancer growth potential indefinitely. The optimal dose is the minimum dose that maintains the tumor burden under a desired limit (e.g., less than 120% of the initial burden, stable disease according to RECIST criteria). To this end, we used the optimal control process described in Subsection **Optimal control**. We first solved the optimality system for a range of different weight values *B* in Equation (8) for all four patients. For each value of *B*, we obtained a time-dependent optimal dose *u*\*. We numerically solved the system for 1460 days (4 years). In this study, we assumed that 4 years is the maximum treatment response owing to computation time. It is worth noting that the median progression-free survival of patients with metastatic melanoma under continuous MTD-targeted therapy ranges from to 11-15 months^53, 54^.

To illustrate this further, we considered the case of Patient 2. The surface plot in Fig 6 A shows the control profiles for different values of *B*. Fig 6 (B) shows the desired tumor burden as a function of *B*. In this study, the desired limit was set as the initial tumor burden. A value of *B* of 691.9 results in containing the tumor at the initial volume (solid blue line in Fig 6 B) and the corresponding time-dependent dose of *u*\* indicated by the solid blue line in 5A (approximately 57% fraction of MTD until almost the end of simulation). This dose could maintain tumor volume under the initial burden (Fig 5 C). The blue line in Fig 5C shows the tumor at the initial volume, and the corresponding dose is shown in Fig 5A by the same-colored lines. The associated volume of the R cells is shown in Fig 5D. An MTD can rapidly decrease the tumor volume by approximately 75 days. However, the tumor volume grows back and relapses by approximately the 168th day and subsequently reaches the carrying capacity. This is due to the growth of R cells (Fig 5D) and extinction of the S cells due to MTD. In optimal dosing, the dose contains the tumor volume at the initial level. The optimal dose modulated the net S-cell growth rate and inter-species competition in favor of the S cells. As a result, R cells die, and the cancer reaches the S-cell-only equilibrium. Because the model was simulated for a period of 4 years, the solution showed a plateau of constant dose at almost all times, followed by a dose reduction before the end of the time interval, which resulted in elevated tumor volume by the end. However, by simulating for a longer period, it can be shown that the cancer can be contained indefinitely with a constant dose at the plateau. We also obtained similar results for the other patients, as shown in Fig 12, 13, and 14 in the Appendix. The optimal dose profiles to contain the tumor at its initial volume are shown in Fig 6 for all patients.

**Figure 5.**
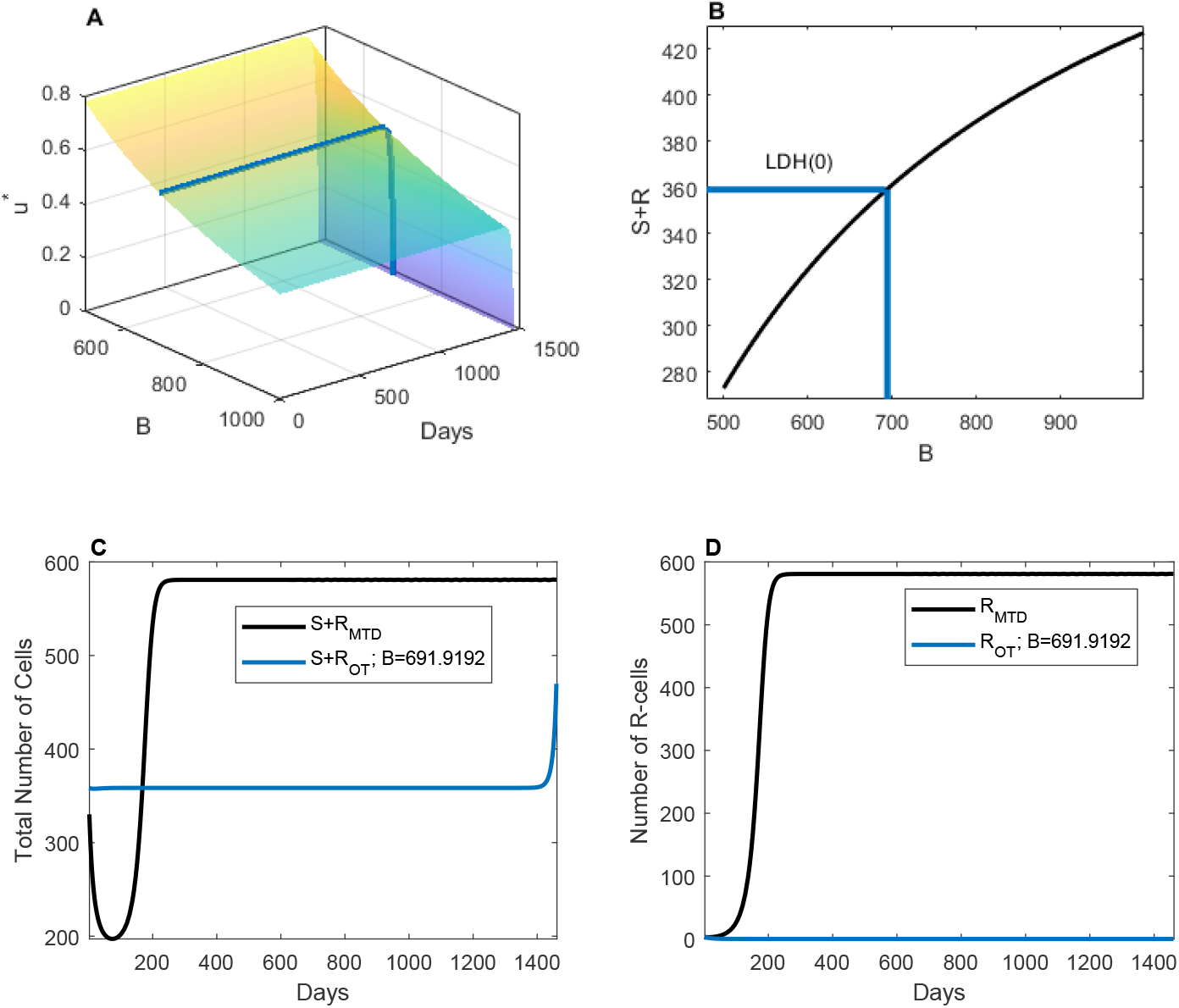
(A) The surface plot shows time-dependent optimal dose for a range of values of *B* for Patient 2. (B) The black line indicates the sustained cancer volume for different values of *B*. The blue line show the value of B for which the cancer volume is contained at the initial volume. Corresponding dose is shown in (A) by the same-colored lines. (C) The blue and black lines show the change in the total cancer volume with optimal dose (contained at the initial volume) and with MTD, respectively. (D) The blue and black lines show the change in the number of R-cells with optimal dose (contained at the initial volume) and with MTD, respectively.

**Figure 6.**
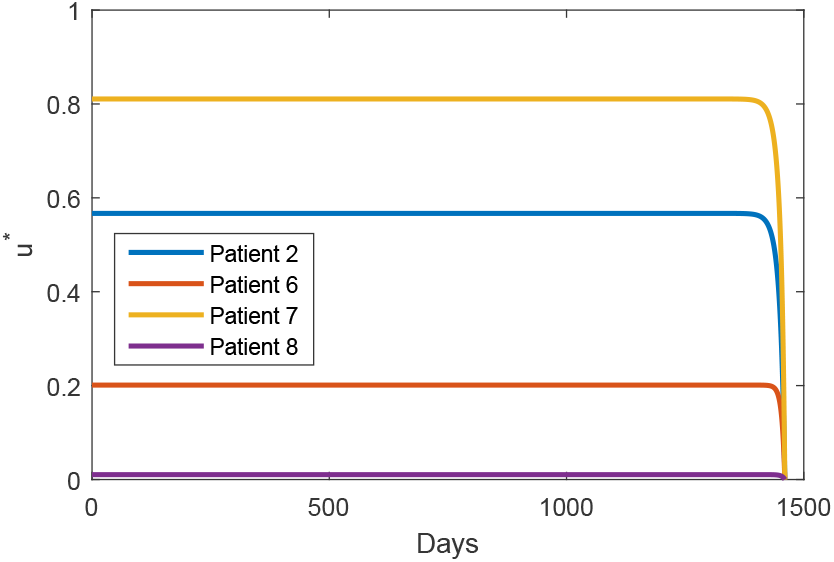
Time-dependent optimal dose has been shown for all the patients. *u*\*(*t*) = 1 corresponds to the MTD. Therefore, in all patients, we observed that a time-dependent dose smaller than the MTD is recommended for optimal therapy.

### Comparing the optimal dose continuous therapy with adaptive therapy

Thus far, we have discussed the effect of continuous therapy with a modulated dose (within the EDW). To compare the treatment outcomes with treatment on and off AT, we simulated AT for all four patients with various dose and pause levels (please find the definition of AT in the section **Adaptive therapy**. To illustrate further, we showed the temporal evolution of tumor burden changes with a fixed pause level of 0.5, and three different doses of 0.5, 0.7, and 0.9, respectively (Fig. 7A). At any of the three different doses, the tumor volume failed to reach the pause level of 0.5, resulting in no treatment vacation. Depending on the dose level, the final tumor volume varied significantly from 85% to approximately 160% of the initial volume. A 50% reduction in dose from MTD (dose 0.5) was not sufficient to reduce tumor burden (Figure 7 an increased blue line), but the dose was able to maintain tumor volume at about 105% of the initial burden. Increasing the dose to 0.7 (70% of the MTD) decreased the tumor burden by approximately 15% from the initial burden (Fig 7, orange line). A further increase to 0.9 reduced the initial burden more rapidly, but later increased it to approximately 160% of the initial volume, consisting of only the R cell (Fig 7, yellow line). If a different pause level of AT is applied to patient 2, maintaining the tumor burden below the initial level can be achieved. For example, an AT with a dose of 0.9 (90% of MTD) and a pause level of 0.65 or 0.7 led to successful tumor burden control (Fig. 7B, yellow and orange lines). A lower pause level (0.6) failed to maintain tumor burden as the burden never reaches 0.6 of the initial with 0.9 of MTD for the patient.

**Figure 7.**
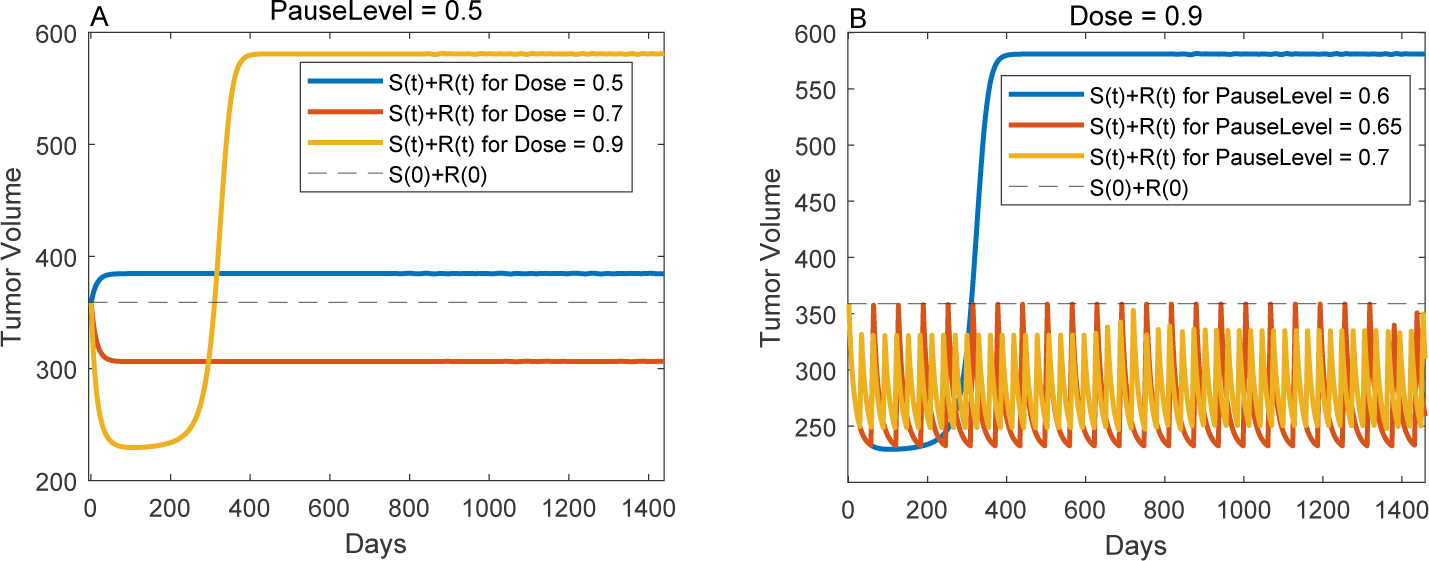
Temporal evolution of tumor burden with various AT strategies. (A) AT with pause level 0.5, dose level 0.5 (blue), 0.7 (orange), and 0.9 (yellow). The gray dashed line indicates the initial tumor burden. (B) AT with a dose level of 0.9 and pause level of 0.6 (blue), 0.65 (orange), and 0.7 (yellow).

We simulated AT for all four patients with various pause levels (50% to 90% of the initial volume) and doses (0 to 100% of MTD), and quantified the TTP for each case. The TTP for each patient is shown in Fig. 8. It is worth noting that TTP is set to 1 if the tumor burden increases from the start of the treatment. Interestingly, a patient-specific dose window ([0.57, 0.84] for patient 2, [0.21, 1] for patient 6, [0.815, 0.82] for patient 7, and [0.015, 0.74] for patient 8 exists, which results in maximum TTP irrespective of the *PauseLevel* (Fig 8 yellow). For doses beyond this window, the pause level can change the TTP. The dose level mostly determines the TTP. For patient 2, if an AT with a dose between 0.57 and 0.84 resulted in the same TTP of 1460 days regardless of the pause level (Fig. 8A, yellow heat map between dose levels of 0.57 and 0.84). A higher dose level required varying pause levels between 0.6 and 0.9 to achieve the same TTP (1460 days). For patient 6, a dose level below 0.21 leads to an increase in tumor volume from treatment initiation (Fig 8B blue shaded region). A dose higher than 0.21 resulted in a TTP of 1460 days regardless of the varying pause level. For patient 7, we observed a good spot in the dose range of 0.815 to 0.82, which led to a TTP of 1460 days (Fig 8C). Interestingly, a dose higher than 0.82 decreased TTP. For Patient 8, our simulations show that a longer TTP could be achieved even with a very low dose (less than 5%) (Fig 8 D). If the dose was increased from 75 to 85%, the pause level was slightly higher than 0.5 for a TTP of 1460 days. Taken together, our AT simulations show that there exists a patient-specific EDW for a large TTP irrespective of the pause level (e.g., 1460 days, approximately four years), which we denote as the EDW (*EDW*_*AT*_).

**Figure 8.**
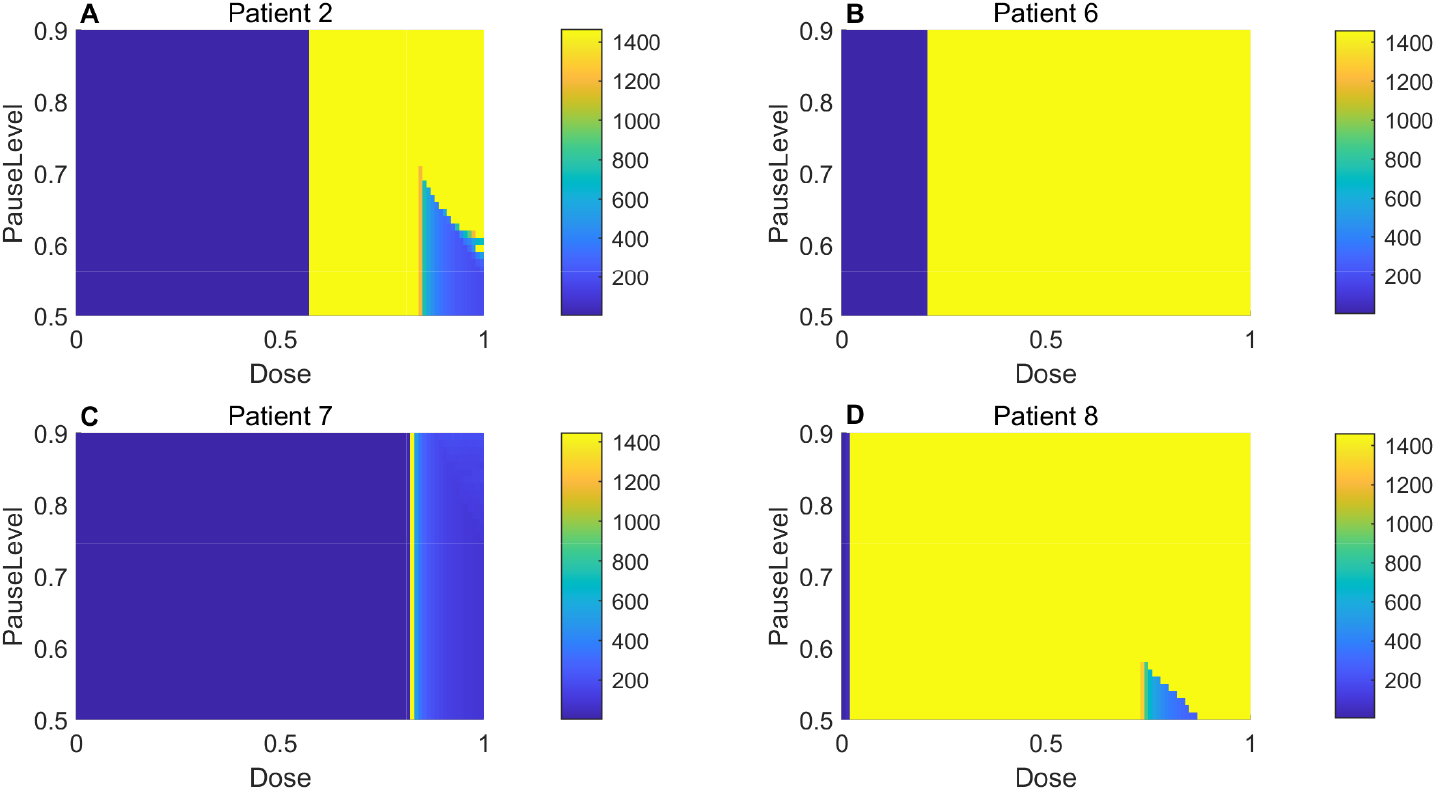
The TTP in days under AT with different pause level and dose for all the patients.

### Defining patient specific MED

In the above section, we learned that there is a dosing interval for each patient in which TTP is maximum and insensitive to pause level. We denote this as the EDW (EDW_*AT*_) for adaptive therapy. To compare the previously derived EDW (shown in columns 6 and 7 of Table 1) with EDW_*AT*_, we normalize the EDW by dividing with the respective value of *δ* because *δ* is the drug-induced rate under MTD therapy and EDW_*AT*_ is a fraction of MTD. We refer it as normalized effective dose window (NEDW). Figure 9 shows a comparison of EDW_*AT*_ (black dotted line) with NEDW (blue dashed line), and the dose at the plateau of the optimal dose profile (asterisk (*)). For all patients, the lower bound of EDW_*AT*_ and NEDW coincided (up to two decimal places) with the optimal dose. The upper bounds of EDW_*AT*_ and NEDW are the same, except for Patient six. Overall, doses belonging to the EDW can extend TTP under both continuous and adaptive treatments. Combining our analysis from three different perspectives, we concluded that the optimal dose that approximately coincides with the lower bounds of EDW_*AT*_ and EDW is the MED.

**Figure 9.**
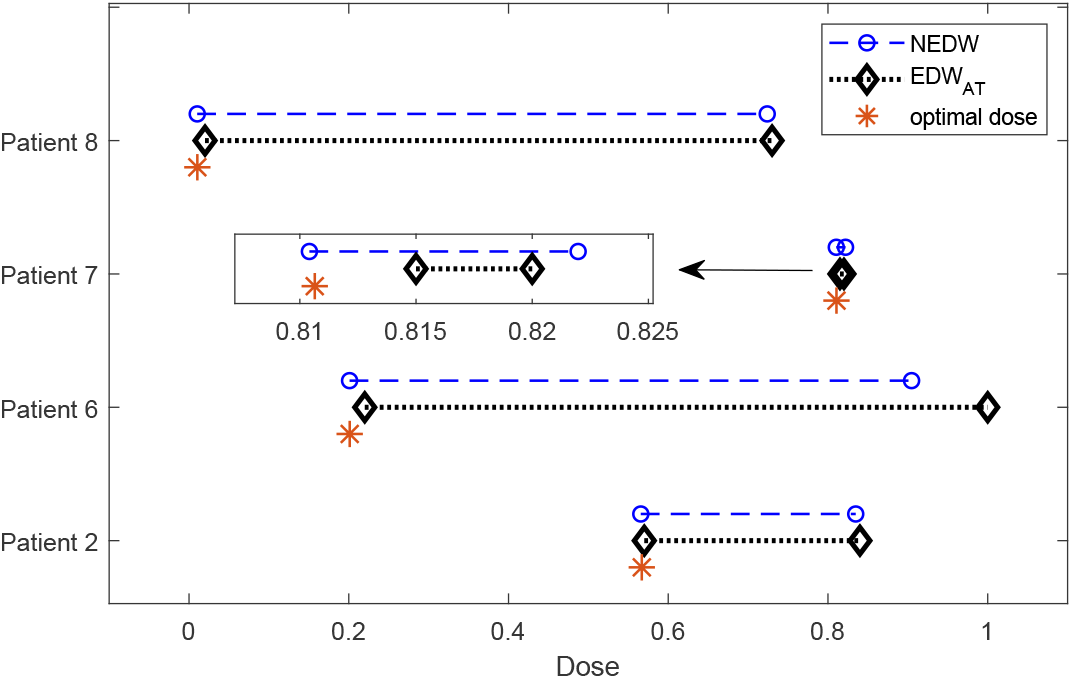
The blue dashed line and black dotted lines resembles the NEDW and EDW_*AT*_. The orange asterisk (*) indicates the optimal dose, designated as the MED. The dose windows for patient seven are very narrow, which is magnified in the inset.

## Discussion and Conclusion

This study investigated whether a sweet spot exists at which the drug-induced death rate holds the tumor growth and suppresses resistance simultaneously. Our analysis of the dynamics of the model identified the EDW (in Equation (13)) and presented the necessary conditions for its existence. To demonstrate this theoretical idea in a real-life scenario, we defined a patient-specific EDW for patients with advanced metastatic melanoma. Further analysis, using the optimum control theory, identified a personalized MED for patients. We compared the treatment outcomes of continuous therapy with an optimized dose using adaptive therapies with varying doses and pause levels. Our simulations show that there exists an EDW (EDW_*AT*_) for which AT offers indefinite control below the initial volume, irrespective of the pause level. Indefinite control has also been reported by Stobl et al. when the tumor cell turnover is high^15^.

Doses smaller than the MTD have been applied in various preclinical and clinical studies. For instance, Mach et al. performed experiments on mouse models of xenograft human pancreatic cancer to determine the efficacy of different doses of liposomal curcumin over a period of four weeks. They demonstrated that a dose of 20*mg/kg* reduces the tumor volume to about 42% of the volume compared to the tumor volume without treatment. Doubling the dose to 40 mg/kg did not lead to a significant reduction in tumor volume. Instead, it has been shown to be a less effective (37% reduction vs. 42% reduction compared with tumor volume without any treatment)^55^. In a mouse model of ovarian cancer, MTD was found to increase the tumor volume by approximately four times, while 20% of the MTD maintains a stable tumor volume of approximately 130% of the initial volume over a period of three weeks^56^.

Our analysis rests on the key assumption that the initial tumor burden of the patient is not immediately life-threatening; rather, it is tolerable. The definition of tolerable tumor volume that is related to the quality of life is obscure^57^, as it is associated with multidimensional factors, such as symptom burden, age, cancer type, and patient expectations. For instance, in a longitudinal study of approximately 500 patients with different types of cancer, patients with stomach, esophageal, hepatobiliary, or head and neck cancer had higher distress levels than other patients^58^. In addition, in phase III trials of prostate cancer, it has been reported that stabilizing the symptom burden is not correlated with the survival rate^59^. However, in most solid cancers, maintaining the sum of the diameters of the target lesions is a treatment response criterion. According to the RECIST 1.1 criteria, a less than 20% increase in the sum is defined as stable disease^48^. A recently reported treatment outcome of AT demonstrated that chronic control of the disease burden could be more effective in improving patient survival^14^. A volume higher than the initial volume is shown to be more effective than aiming at rapid reduction of tumor volume with MTD therapy in theoretical (110% of the initial volume^2^) as well as preclinical studies (125% of the initial volume^19^).

In developing mathematical models for clinical applications, it is challenging to quantify the relevant parameters with limited data. Several sets of parameter values may produce a similar fit to the data, which leads to the unidentifiability of the parameters and limitations in the model-based quantitative decision-making. Unidentifiability may appear because of structural or practical unidentifiability. While the former requires remodeling, the latter requires more frequent and less noisy data, as shown in Fig 10. In this study, our model was structurally identifiable, but the biomarker data of four of the patients could not produce practically identifiable parameter estimates, which were excluded from further analysis. In such cases, more frequent and less noisy data can result in identifiable parameters. With the identified parameters, we were able to infer tumor dynamics properly and propose a personalized MED therapy. We argue that the MED derived from our approach could lead to increased patient survival and less toxicity. We believe this approach partially supports the notion of a digital twin in cancer care^60, 61^.

**Figure 10.**
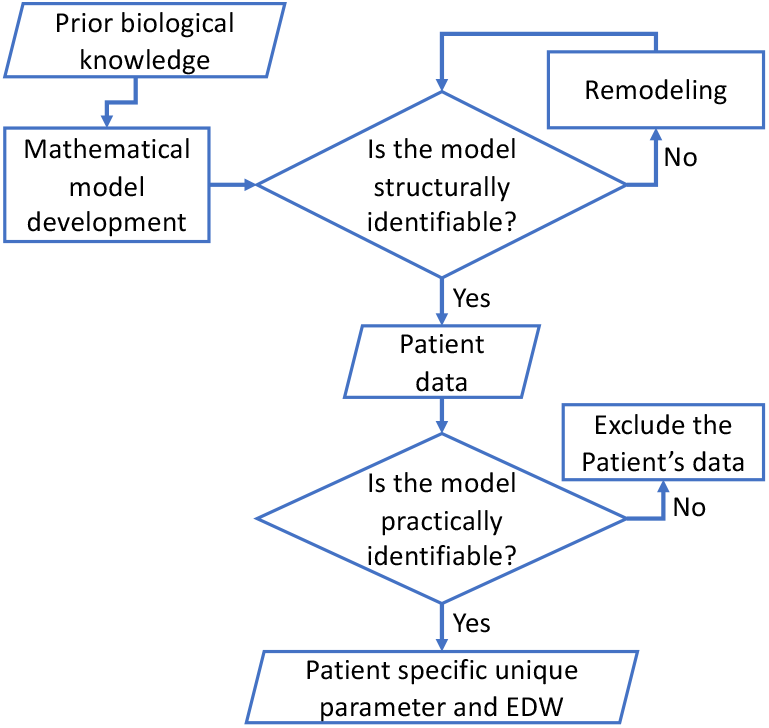
The flowchart illustrates the process of modeling tumor dynamics and estimation of patient specific parameters.

Our model is an abstract representation of tumors in a patient. We assumed that the tumor cell population was a homogeneous mixture of two genetically fixed drug-sensitive and drug-resistant cell populations. In real tumors, cancer cells can be phonetically plastic and a drug can induce resistance^18, 62, 63^. In addition, cancer populations are not well-mixed, but rather spatially organized^64–67^, which can be modulated by heterogeneous tumor microenvironmental factors^68, 69^. Currently, the clinical assessment of treatment response is often performed by analyzing non-spatial tumor burden data (blood level of tumor burden). Therefore, although a spatial model may better represent the tumor, it would be more complex with additional assumptions that cannot be supported by blood biomarkers only.

In summary, our analysis of tumor dynamics identifies the necessary conditions for the existence of an EDW, where its lower bound corresponds to the MED derived by applying optimal control theory. The application of our approach to patients with advanced melanoma identified the personalized MED. Here, we identify such a dose by performing identifiability analysis and calibrating the model to each patient’s tumor burden dynamic data. This study highlights the potential benefits of using mathematical models in clinics by supporting personalized MED decisions. This mathematical model-integrated treatment decision paradigm is crucial for personalized medicine because it facilitates therapy dosing. Therefore, we advocate integrating multiple principles, including predictive mathematical models, to develop novel therapeutic strategies.

## A Appendix

### Mathematical Analysis of the model

#### Theorem A.1.

*The trivial equilibrium* (0, 0) *is unstable*.

*Proof*. Jacobian of system (1) at the trivial equilibrium (*J*_0_),

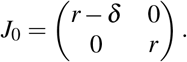

*J*_0_ is a diagonal matrix with a positive eigenvalue *r*. Therefore, trivial equilibrium is unstable. □

#### Theorem A.2.

*The S-only equilibrium exists if δ < r, and is locally asymptotically stable if* 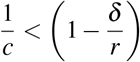,

*Proof*. The S-only equilibrium is given by 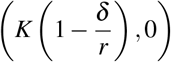: Because we address the non-negative number of cell populations, this equilibrium exists for *δ < r*. Moreover, by evaluating the Jacobian of the system (1) at the S-only equilibrium (*J*_*S*_),

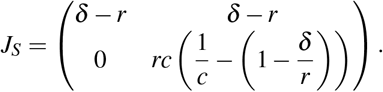

*J*_*S*_ is an upper triangular matrix. Therefore, it is sufficient to show that the diagonal elements are negative to prove local asymptotic stability. According to the requirement for the existence of an S-only equilibrium, the first diagonal element *δ* − *r <* 0. The second diagonal element is negative, when 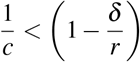. This completes the proof. □

#### Theorem A.3.

*R-only always exists and is locally asymptotically stable*.

*Proof*. The R-only equilibrium is given by (0, *K*) and exists for any positive *K*. The Jacobian of the system (1) at R-only equilibrium (*J*_*R*_) is given by

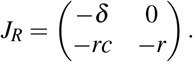

As, it is a diagonal matrix with all diagonal elements being negative, the R-only equilibrium is locally asymptotically stable. □

#### Theorem A.4.

*Co-existence equilibrium exists if* 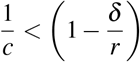 *and is unstable when exists*.

*Proof*. The coexistence equilibrium holds for the expressions 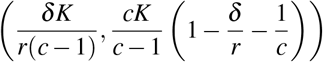. This is positive if *c >* 1 and 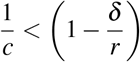. The Jacobian at the coexistence equilibrium upon simplification reduces to the following form:

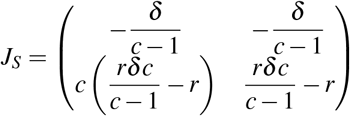

The determinant of the above matrix 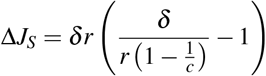 is negative if 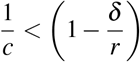. Therefore, it is unstable when exists. □

### B Profile likelihood of patients with practically non-identifiable parameters

**Figure 11.**
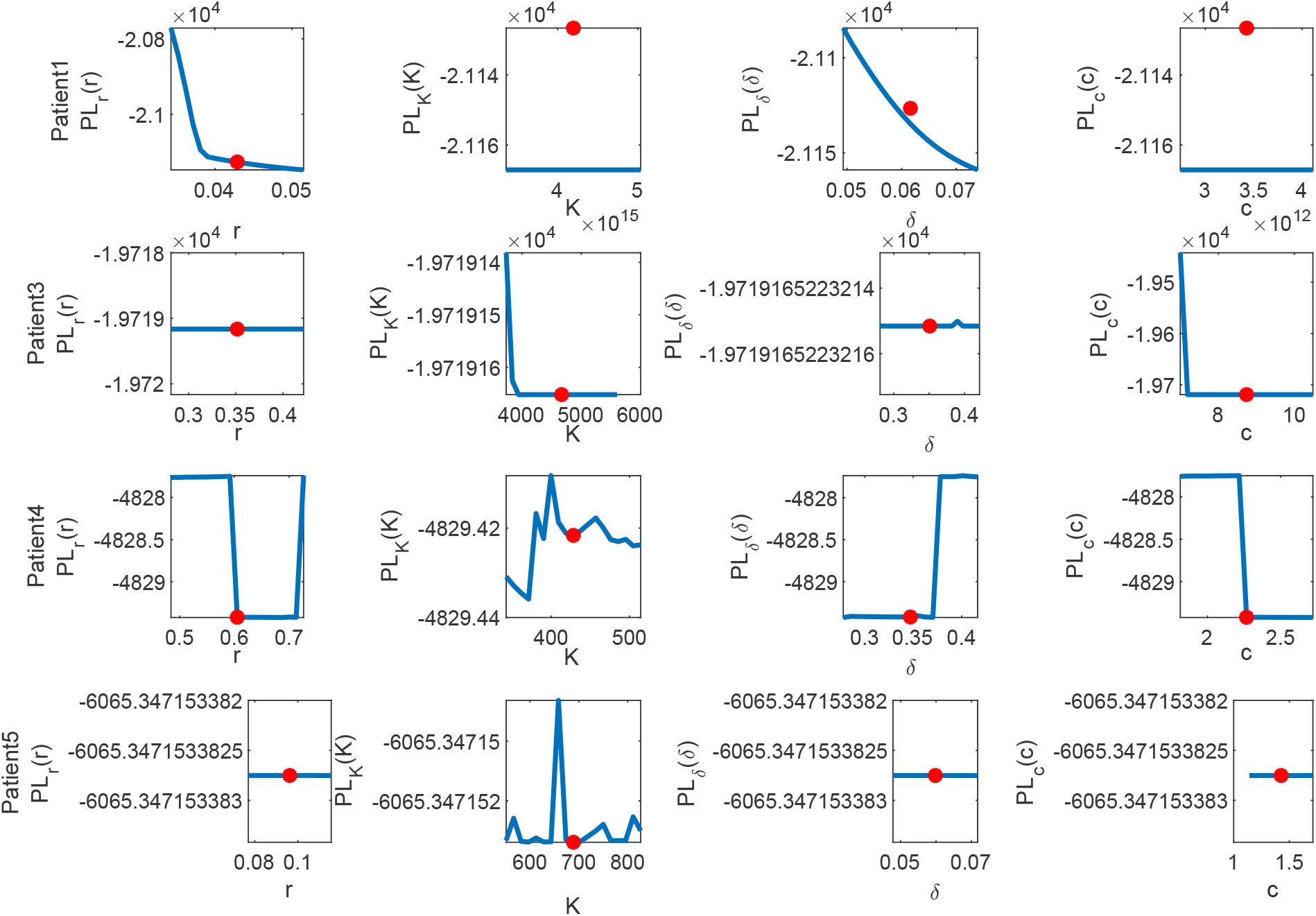
Each row shows the profile likelihood for each of the parameters for Patient 1, 3, 4 and 5 respectively, where y axis is 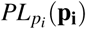 defined in equation (6) and x-axis is the perturbed parameter *p*_*i*_ ∈ {*r, K, δ, c*}. The profile likelihoods show that the estimates do not minimize the likelihood, and are hence unidentifiable.

### C. Optimal dose

#### Theorem C.1.

*Provided that there exists an optimal dose u*\*(*t*) *corresponding to S*\*(*t*) *and R*\*(*t*) *that minimizes the cost function* (8) *subject to* (7), *there exist adjoint variables λ*_1_ *and λ*_2_ *given by* (10) *with transversality conditions λ*_1_(*T*) = 0 *and λ*_2_(*T*) = 0. *Moreover, the optimal dose, u*\*(*t*), *is given by* (11).

*Proof*. Pontryagin’s maximum principle^46, 47^ converts the optimization problem into a point-wise minimization of the Hamiltonian ℋ ((9)). The adjoint system is deduced from the following derivatives:

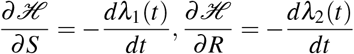

The optimality condition 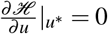 yields optimal dose *u*\*(*t*). Within the bound 0 ≤ *u*(*t*) ≤ 1, this is given by (11). □

#### C.1 Numerical solution of the optimality system

**Figure 12.**
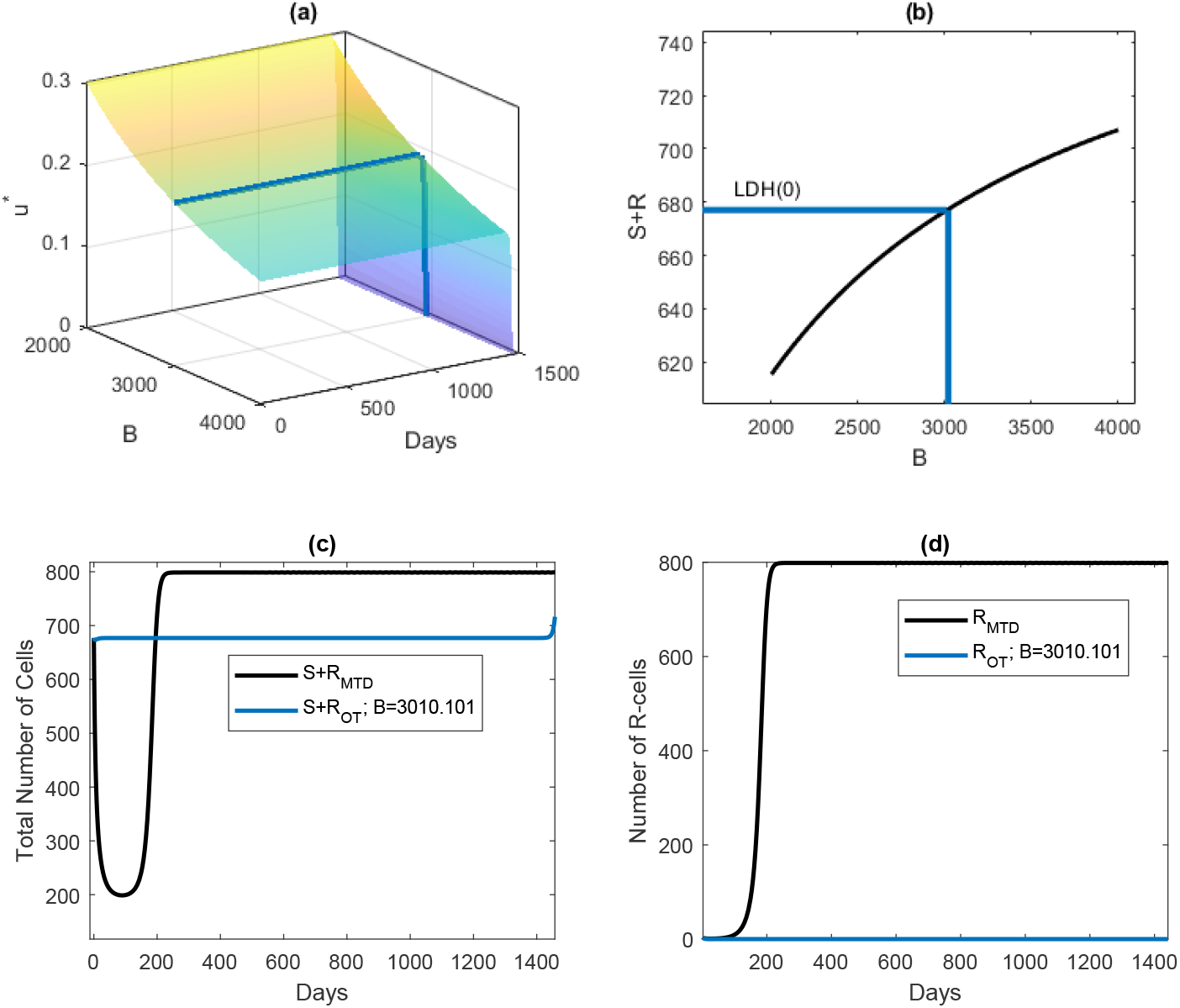
(a) The surface plot shows time-dependent optimal dose for a range of values of *B* for Patient 6. (b) The black line indicates the sustained cancer volume for different values of *B*. The blue line show the value of B for which the cancer volume is contained at the initial volume. Corresponding dose is shown in (a) by the same-colored lines. (c) The blue and black lines show the change in the total cancer volume with optimal dose (contained at the initial volume) and with MTD, respectively. (d) The blue and black lines show the change in the number of R-cells with optimal dose (contained at the initial volume) and with MTD, respectively

**Figure 13.**
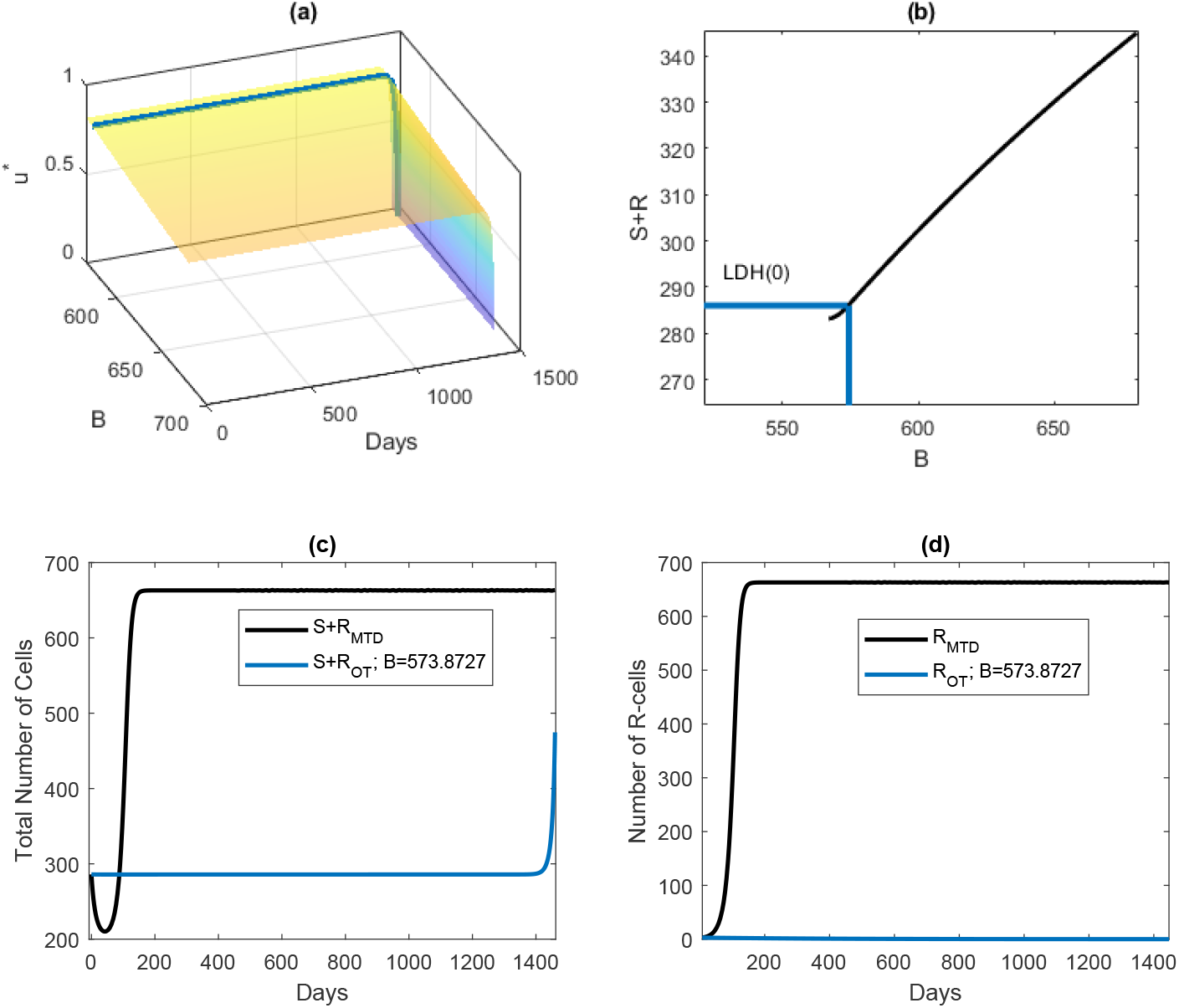
(a) The surface plot shows time-dependent optimal dose for a range of values of *B* for Patient 7. (b) The black line indicates the sustained cancer volume for different values of *B*. The blue line show the value of B for which the cancer volume is contained at the initial volume. Corresponding dose is shown in (a) by the same-colored lines. (c) The blue and black lines show the change in the total cancer volume with optimal dose (contained at the initial volume) and with MTD, respectively. (d) The blue and black lines show the change in the number of R-cells with optimal dose (contained at the initial volume) and with MTD, respectively

**Figure 14.**
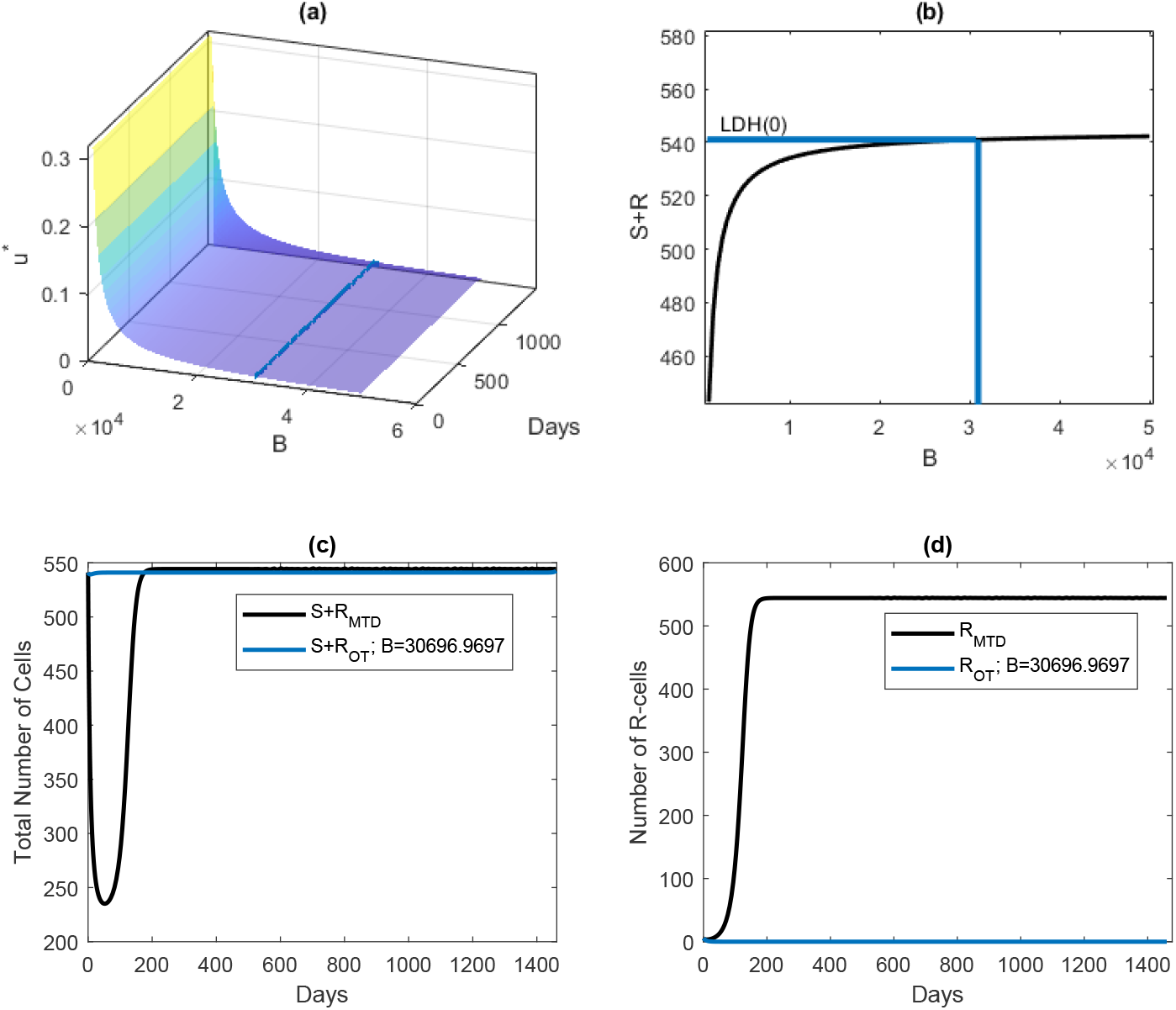
(a) The surface plot shows time-dependent optimal dose for a range of values of *B* for Patient 8. (b) The black line indicates the sustained cancer volume for different values of *B*. The blue line show the value of B for which the cancer volume is contained at the initial volume. Corresponding dose is shown in (a) by the same-colored lines. (c) The blue and black lines show the change in the total cancer volume with optimal dose (contained at the initial volume) and with MTD, respectively. (d) The blue and black lines show the change in the number of R-cells with optimal dose (contained at the initial volume) and with MTD, respectively.

## Author contributions statement

Conceptualization: Masud M A, Eunjung Kim; Data curation: Masud M A, Eunjung Kim; Formal analysis: Masud M A; Funding acquisition: Eunjung Kim; Investigation: Masud M A, Jae-Young Kim, Eunjung Kim; Methodology: Masud M A; Supervision: Jae-Young Kim, Eunjung Kim; Visualization: Masud M A; Writing – original draft: Masud M A, Jae-Young Kim, Eunjung Kim; Writing – review and editing: Jae-Young Kim and Eunjung Kim. All authors have reviewed the manuscript.

## References

1. Le Tourneau, C., Lee, J. J. & Siu, L. L. Dose Escalation Methods in Phase I Cancer Clinical Trials. JNCI: J. National Cancer Inst. 101, 708–720, DOI: 10.1093/jnci/djp079 (2009).

2. Gallaher, J. A., Enriquez-Navas, P. M., Luddy, K. A., Gatenby, R. A. & Anderson, A. R. Spatial heterogeneity and evolutionary dynamics modulate time to recurrence in continuous and adaptive cancer therapies. Cancer Research 78, 2127–2139, DOI: 10.1158/0008-5472.CAN-17-2649 (2018).

3. Smalley, I. et al. Leveraging transcriptional dynamics to improve braf inhibitor responses in melanoma. EBioMedicine 48, 178–190, DOI: https://doi.org/10.1016/j.ebiom.2019.09.023 (2019).

4. Jensen, N. F. et al. Establishment and characterization of models of chemotherapy resistance in colorectal cancer: Towards a predictive signature of chemoresistance. Mol. Oncol. 9, 1169–1185, DOI: https://doi.org/10.1016/j.molonc.2015.02.008 (2015).

5. Carrère, C. Optimization of an in vitro chemotherapy to avoid resistant tumours. J. Theor. Biol. 413, 24–33, DOI: https://doi.org/10.1016/j.jtbi.2016.11.009 (2017).

6. Bacevic, K. et al. Spatial competition constrains resistance to targeted cancer therapy. Nat. communications 8, DOI: 10.1038/s41467-017-01516-1 (2017).

7. Rozados, V. et al. Metronomic therapy with cyclophosphamide induces rat lymphoma and sarcoma regression, and is devoid of toxicity. Annals Oncol. 15, 1543–1550, DOI: https://doi.org/10.1093/annonc/mdh384 (2004).

8. Kareva, I., Waxman, D. J. & Lakka Klement, G. Metronomic chemotherapy: An attractive alternative to maximum tolerated dose therapy that can activate anti-tumor immunity and minimize therapeutic resistance. Cancer Lett. 358, 100–106, DOI: https://doi.org/10.1016/j.canlet.2014.12.039 (2015).

9. Sachs, J. R., Mayawala, K., Gadamsetty, S., Kang, S. P. & de Alwis, D. P. Optimal dosing for targeted therapies in oncology: Drug development cases leading by example. Clin. Cancer Research 22, 1318–1324, DOI: 10.1158/1078-0432.CCR-15-1295 (2016). https://clincancerres.aacrjournals.org/content/22/6/1318.full.pdf.

10. Corbaux, P. et al. Clinical efficacy of the optimal biological dose in early-phase trials of anti-cancer targeted therapies. Eur. J. Cancer 120, 40–46, DOI: https://doi.org/10.1016/j.ejca.2019.08.002 (2019).

11. Gasparini, G. Metronomic scheduling: the future of chemotherapy? The Lancet. Oncol. 2 12, 733–740 (2001).

12. Shu, Y., Weng, S. & Zheng, S. Metronomic chemotherapy in non-small cell lung cancer. Oncol. letters 20 6, 307 (2020).

13. Gatenby, R. A., Silva, A. S., Gillies, R. J. & Frieden, B. R. Adaptive therapy. Cancer Research 69, 4894–4903, DOI: 10.1158/0008-5472.CAN-08-3658 (2009).

14. Zhang, J., Cunningham, J. J., Brown, J. S. & Gatenby, R. A. Integrating evolutionary dynamics into treatment of metastatic castrate-resistant prostate cancer. Nat. Commun. 8 (2017).

15. Strobl, M. A. et al. Turnover modulates the need for a cost of resistance in adaptive therapy. Cancer Research 81, 1135–1147, DOI: 10.1158/0008-5472.CAN-20-0806 (2021).

16. Strobl, M. A. R. et al. Spatial structure impacts adaptive therapy by shaping intra-tumoral competition. bioRxiv DOI: 10.1101/2020.11.03.365163 (2021).

17. Brady-Nicholls, R. et al. Predicting patient-specific response to adaptive therapy in metastatic castration-resistant prostate cancer using prostate-specific antigen dynamics. Neoplasia 23, 851–858, DOI: https://doi.org/10.1016/j.neo.2021.06.013 (2021).

18. Kim, E., Brown, J. S., Eroglu, Z. & Anderson, A. R. Adaptive therapy for metastatic melanoma: Predictions from patient calibrated mathematical models. Cancers 13 (2021).

19. Enriquez-Navas, P. M. et al. Exploiting evolutionary principles to prolong tumor control in preclinical models of breast cancer. Sci. Transl. Medicine 8, 327ra24–327ra24, DOI: 10.1126/scitranslmed.aad7842 (2016).

20. Viossat, Y. & Noble, R. A theoretical analysis of tumour containment. Nat. Ecol. Evol. 5, 826–83 (2021).

21. Hansen, E. & Read, A. F. Modifying adaptive therapy to enhance competitive suppression. Cancers 12, DOI: 10.3390/cancers12123556 (2020).

22. Gatenby, R. A change of strategy in the war on cancer. Nature 459, 508–509 (2009).

23. Martin, R. B., Fisher, M. E., Minchin, R. F. & Teo, K. L. Optimal control of tumor size used to maximize survival time when cells are resistant to chemotherapy. Math. biosciences 110 2, 201–19 (1992).

24. Pouchol, C., Clairambault, J., Lorz, A. & Trélat, E. Asymptotic analysis and optimal control of an integro-differential system modelling healthy and cancer cells exposed to chemotherapy. J. de Math. Pures et Appliquées 116, 268–308, DOI: https://doi.org/10.1016/j.matpur.2017.10.007 (2018).

25. Almeida, L., Bagnerini, P., Fabrini, G., Hughes, B. D. & Lorenzi, T. Evolution of cancer cell populations under cytotoxic therapy and treatment optimisation: insight from a phenotype-structured model. ESAIM: Math. Model. Numer. Analysis 53, 1157–1190, DOI: 10.1051/m2an/2019010 (2019).

26. Cunningham, J. J., Brown, J. S., Gatenby, R. A. & Stan?ková, K. Optimal control to develop therapeutic strategies for metastatic castrate resistant prostate cancer. J. Theor. Biol. 459, 67–78, DOI: https://doi.org/10.1016/j.jtbi.2018.09.022 (2018).

27. Cunningham, J. et al. Optimal control to reach eco-evolutionary stability in metastatic castrate-resistant prostate cancer. PLOS ONE 15, 1–24, DOI: 10.1371/journal.pone.0243386 (2020).

28. Ledzewicz, U. et al. On drug resistance and metronomic chemotherapy: A mathematical modeling and optimal control approach. Math. Biosci. Eng. 14, 217–235, DOI: 10.3934/mbe.2017014 (2017).

29. Bondarenko, M. et al. Metronomic chemotherapy modulates clonal interactions to prevent drug resistance in non-small cell lung cancer. Cancers 13, DOI: 10.3390/cancers13092239 (2021).

30. Marquis, A. D., Arnold, A., Dean-Bernhoft, C., Carlson, B. E. & Olufsen, M. S. Practical identifiability and uncertainty quantification of a pulsatile cardiovascular model. Math. Biosci. 304, 9–24, DOI: https://doi.org/10.1016/j.mbs.2018.07.001 (2018).

31. Eisenberg, M. C. & Hayashi, M. A. Determining identifiable parameter combinations using subset profiling. Math. Biosci. 256, 116–126, DOI: https://doi.org/10.1016/j.mbs.2014.08.008 (2014).

32. Olufsen, M. S. & Ottesen, J. T. A practical approach to parameter estimation applied to model predicting heart rate regulation. J. Math. Biol. 67, 39–68, DOI: 10.1007/s00285-012-0535-8 (2013).

33. Meshkat, N., Kuo, C.E.-z. & DiStefano, J., III. On finding and using identifiable parameter combinations in nonlinear dynamic systems biology models and combos: A novel web implementation. PLOS ONE 9, 1–14, DOI: 10.1371/journal.pone.0110261 (2014).

34. Meshkat, N., Eisenberg, M. & DiStefano, J. J. An algorithm for finding globally identifiable parameter combinations of nonlinear ode models using Grobner bases. Math. Biosci. 222, 61–72, DOI: https://doi.org/10.1016/j.mbs.2009.08.010 (2009).

35. Meshkat, N., Anderson, C. & DiStefano, J. J. Finding identifiable parameter combinations in nonlinear ode models and the rational reparameterization of their input–output equations. Math. Biosci. 233, 19–31, DOI: https://doi.org/10.1016/j.mbs.2011.06.001 (2011).

36. Meshkat, N., Anderson, C. & DiStefano III, J. J. Alternative to ritt’s pseudodivision for finding the input-output equations of multi-output models. Math. Biosci. 239, 117–123, DOI: https://doi.org/10.1016/j.mbs.2012.04.008 (2012).

37. Ljung, L. & Glad, T. On global identifiability for arbitrary model parametrizations. Automatica 30, 265–276, DOI: https://doi.org/10.1016/0005-1098(94)90029-9 (1994).

38. Bellu, G., Saccomani, M. P., Audoly, S. & D’Angiò, L. Daisy: A new software tool to test global identifiability of biological and physiological systems. Comput. Methods Programs Biomed. 88, 52–61, DOI: https://doi.org/10.1016/j.cmpb.2007.07.002 (2007).

39. Scarpiniti, M. & Villaverde, A. F. Observability and structural identifiability of nonlinear biological systems. Complexity 2019, DOI: 10.1155/2019/8497093 (2019).

40. MAM.Islam, M. H., Mamun, K. A., Kim, B. N. & Kim, S. Covid-19 transmission: Bangladesh perspective. Mathematics 8, DOI: 10.3390/math8101793 (2020).

41. Lira-Parada, P. A., Pettersen, E., Biegler, L. T. & Bar, N. Implications of dimensional analysis in bioreactor models: Parameter estimation and identifiability. Chem. Eng. J. 417, 129220, DOI: https://doi.org/10.1016/j.cej.2021.129220 (2021).

42. Cobelli, C. & DiStefano, J. J. Parameter and structural identifiability concepts and ambiguities: a critical review and analysis. Am. J. Physiol. Integr. Comp. Physiol. 239, R7–R24, DOI: 10.1152/ajpregu.1980.239.1.R7 (1980).

43. Rothenberg, T. J. Identification in parametric models. Econometrica 39, 577–591 (1971).

44. Murphy, S. A. & Van Der Vaart, A. W. On profile likelihood. J. Am. Stat. Assoc. 95, 449–465, DOI: 10.1080/01621459.2000.10474219 (2000).

45. Fleming, W. H. & Rishel, R. W. Deterministic and Stochastic Optimal Control (Springer Verlag, 1975).

46. Pontryagin, L. S., Boltyanskii, V. G., Gamkrelidze, R. V. & Mischenko, E. F. The Mathematical Theory of Optimal Processes (Wiley, New Jersey, 1962).

47. Lenhart, S. & Workman, J. T. Optimal Control Applied to Biological Models (Chapman and Hall CRC, London, 2007).

48. Eisenhauer, E. et al. New response evaluation criteria in solid tumours: Revised recist guideline (version 1.1). Eur. J. Cancer 45, 228–247, DOI: https://doi.org/10.1016/j.ejca.2008.10.026 (2009).

49. Weinstein, D., Leininger, J., Hamby, C. & Safai, B. Diagnostic and prognostic biomarkers in melanoma. The J. clinical aesthetic dermatology 7 (2014).

50. Gershenwald, J. E. et al. Melanoma staging: Evidence-based changes in the american joint committee on cancer eighth edition cancer staging manual. CA: A Cancer J. for Clin. 67, 472–492, DOI: https://doi.org/10.3322/caac.21409 (2017).

51. Bozic, I. & Nowak, M. A. Timing and heterogeneity of mutations associated with drug resistance in metastatic cancers. Proc. National Acad. Sci. 111, 15964–15968, DOI: 10.1073/pnas.1412075111 (2014).

52. Grassberger, C. et al. Patient-specific tumor growth trajectories determine persistent and resistant cancer cell populations during treatment with targeted therapies. Cancer Research 79, 3776–3788, DOI: 10.1158/0008-5472.CAN-18-3652 (2019).

53. Ascierto, P. A. et al. Cobimetinib combined with vemurafenib in advanced brafv600-mutant melanoma (cobrim): updated efficacy results from a randomised, double-blind, phase 3 trial. The Lancet Oncol. 17, 1248–1260, DOI: https://doi.org/10.1016/S1470-2045(16)30122-X (2016).

54. Dummer, R. et al. Encorafenib plus binimetinib versus vemurafenib or encorafenib in patients with braf-mutant melanoma (columbus): a multicentre, open-label, randomised phase 3 trial. The Lancet Oncol. 19, 603–615, DOI: https://doi.org/10.1016/S1470-2045(18)30142-6 (2018).

55. Mach, C. M., Mathew, L., Mosley, S. A., Kurzrock, R. & Smith, J. A. Determination of minimum effective dose and optimal dosing schedule for liposomal curcumin in a xenograft human pancreatic cancer model. Anticancer Research 29, 1895–1899 (2009). https://ar.iiarjournals.org/content/29/6/1895.full.pdf.

56. Caumanns, J. J. et al. Low-dose triple drug combination targeting the pi3k/akt/mtor pathway and the mapk pathway is an effective approach in ovarian clear cell carcinoma. Cancer Lett. 461, 102–111, DOI: https://doi.org/10.1016/j.canlet.2019.07.004 (2019).

57. Bottomley, A. The cancer patient and quality of life. The Oncol. 7, 120–125, DOI: 10.1634/theoncologist.7-2-120 (2002).

58. Vogt, J. et al. Symptom burden and palliative care needs of patients with incurable cancer at diagnosis and during the disease course. The Oncol. 26, e1058–e1065, DOI: 10.1002/onco.13751 (2021).

59. Adamowicz, K. Assessment of quality of life in advanced, metastatic prostate cancer: an overview of randomized phase iii trials. Qual. Life Research 26, 813–822, DOI: 10.1007/s11136-016-1429-9 (2017).

60. Hernandez-Boussard, T. et al. Digital twins for predictive oncology will be a paradigm shift for precision cancer care. Nat. Medicine 27, 2065–2066, DOI: 10.1038/s41591-021-01558-5 (2021).

61. Thiong’o, G. M. & Rutka, J. T. Digital twin technology: The future of predicting neurological complications of pediatric cancers and their treatment. Front. Oncol. 11, DOI: 10.3389/fonc.2021.781499 (2022).

62. Greene, J. M., Gevertz, J. L. & Sontag, E. D. Mathematical approach to differentiate spontaneous and induced evolution to drug resistance during cancer treatment. JCO Clin. Cancer Informatics 1–20, DOI: 10.1200/CCI.18.00087 (2019).

63. Greene, J. M., Sanchez-Tapia, C. & Sontag, E. D. Mathematical details on a cancer resistance model. Front. Bioeng. Biotechnol. 8, DOI: 10.3389/fbioe.2020.00501 (2020).

64. Gatenby, R. A., Grove, O. & Gillies, R. J. Quantitative imaging in cancer evolution and ecology. Radiology 269, 8–14, DOI: 10.1148/radiol.13122697 (2013).

65. Gerlinger, M. et al. Intratumor heterogeneity and branched evolution revealed by multiregion sequencing. New Engl. J. Medicine 366, 883–892, DOI: 10.1056/NEJMoa1113205 (2012).

66. Gillies, R., Anderson, A. R. A., Gatenby, R. A. & Morse, D. L. The biology underlying molecular imaging in oncology: from genome to anatome and back again. Clin. radiology 65 7, 517–21 (2010).

67. O’Connor, J. P. et al. Imaging intratumor heterogeneity: Role in therapy response, resistance, and clinical outcome. Clin. Cancer Research 21, 249–257, DOI: 10.1158/1078-0432.CCR-14-0990 (2015).

68. Mumenthaler, S. M. et al. The impact of microenvironmental heterogeneity on the evolution of drug resistance in cancer cells. Cancer Informatics 14, 19 – 31 (2015).

69. Kaznatcheev, A., Peacock, J., Basanta, D., Marusyk, A. & Scott, J. G. Fibroblasts and alectinib switch the evolutionary games played by non-small cell lung cancer. Nat. ecology & evolution 3, 450 – 456 (2019).

